# Bacterial growth, communication and guided chemotaxis in 3D bioprinted hydrogel environments

**DOI:** 10.1101/2021.08.10.455848

**Authors:** Julia Müller, Anna C. Jäkel, Jonathan Richter, Markus Eder, Elisabeth Falgenhauer, Friedrich C. Simmel, F. C. Simmel TU München

## Abstract

Bioprinting of engineered bacteria is of great interest for applications of synthetic biology in the context of living biomaterials, but so far only few viable approaches are available for the printing of gels hosting live *Escherichia coli* bacteria. Here we develop a gentle bioprinting method based on an alginate/agarose bioink that enables precise printing of *E.coli* into three-dimensional hydrogel structures up to 10 mm in height. Addition of a calcium peroxide-based oxygen generation system enables maturation of fluorescent proteins deep within the printed structures. We utilize spatial patterning with the bioprinter to control different types of chemical interaction between bacteria. We first show quorum sensing-based chemical communication between engineered sender and receiver bacteria placed at different positions inside the bioprint, and then demonstrate the fabrication of barrier structures defined by non-motile bacteria that can guide the movement of chemotactic bacteria inside a gel.

## 1. Introduction

Additive manufacturing of biocompatible scaffold structures and 3D printing of gel-encapsulated mammalian cells have become popular approaches for the generation of spatially differentiated cell cultures and models of living tissues, which promise to have a wide range of biomedical applications ^[1]^. Bioprinting of bacterial systems is comparatively less developed, but is expected to be of similar importance for the engineering of spatially organized bacterial communities, which will ultimately lead to the realization of living, bacteria-based biomaterials ^[2]^. Spatial arrangement and confinement of bacterial communities also allows the study and control of their interactions with their environment or other communities under well-defined experimental conditions. This is of particular interest for applications in synthetic biology, which involve spatially distributed synthetic cell-cell communication and division of labor between different types of genetically engineered bacteria.

Functional biomaterials with biosensing ^[3]^ or biocatalytic capabilities ^[4]^ have been previously realized by immobilizing bacteria within various hydrogel-based support matrices, which in some cases also allowed a rough positioning of the bacteria in 3D. At a more macroscopic scale, hydrogel matrices were loaded with bacterial cultures to enable printing of biofilms for the degradation of pollutants ^[5]^, the realization of self-healing structures ^[2b]^, and even bacterial electrodes ^[6]^. More recently, 3D bioprinting techniques were utilized to achieve better spatial control over bacteria-laden structures. Laser-based lithography was utilized to contain bacterial cells within picoliter-sized gel compartments, which enabled the observation of cell-cell interactions and bacterial signaling between small populations ^[7]^. Bacteria were also encapsulated in a Pluronic F127 diacrylate-containing bioink followed by UV-crosslinking, which was used for the realization of printable bacteria-based biosensors ^[8]^. In a different approach,^[2a]^ printed *B. subtilis* spores were mixed with liquid agarose that started new bacterial colonies upon solidification of the gel and incubation after printing. The use of bacterial spores allowed printing at temperatures as a high as 70°C, which would not have been possible with living bacteria. An alternative approach^[9]^ comprises extrusion of homogeneous *E*.*coli* biofilms using a custom-built 3D printer that is loaded with an alginate ink, which solidifies upon contact with a calcium-containing substrate. While this approach does not require high temperatures, only a few layers can be printed as diffusion of calcium ions is required for the solidification of the alginate gel. The majority of these examples resulted in homogeneous bacterial structures containing only a single bacterial clone.

Printing of living cells generally requires a compromise between printability and biocompatibility of the materials used for printing. Most approaches employed to print functional bacterial gels involve a curing step after printing, either using a chemical crosslinker ^[2b, 6, 9]^ or photo-crosslinking ^[7-8]^. The hydrogels used to encapsulate and print the bacteria had to be chosen specifically ^[5]^ to facilitate printing at sufficiently low temperatures, which ensures cell viability and prevents sporulation of spore-forming bacteria.

Previous approaches were not capable of printing live *E*.*coli* bacteria into genuine 3D hydrogel structures, within which the bacteria would execute engineered genetic functions. Such a capability is of major interest, however, as *E*.*coli* is the most developed genetic “workhorse” with a broad range of tools available for engineering biology.

To address this need, in the present work we developed a gentle approach to print living *E*.*coli* bacteria embedded in a non-toxic hydrogel environment composed of a mixture of alginate and agarose, which allowed printing at moderate temperatures and without the need for chemical crosslinking. Addition of calcium peroxide and the enzyme catalase ^[10]^ to the bioink ensured a continuous supply of the bacteria with oxygen within the printed gel over a time course of at least 24 hours.

Using this method, we were able to sequentially print three different bacterial clones in a single layer and express fluorescent proteins at the respective locations. Dual extrusion of bacteria-free and bacteria-laden hydrogels enabled printing of bacteria at programmed positions within 3D structures of heights of up to 10 mm.

Bacterial bioprinting opens up the possibility to precisely define the boundary and initial conditions of dynamical systems composed of bacteria with genetically programmed interactions ^[11]^, which potentially allows to combine the advantages of top-down patterning with self-organized spatiotemporal differentiation for the realization of living biomaterials. To demonstrate this capability, we applied our bioprinting approach to control quorum-sensing based bacterial communication between engineered sender and receiver bacteria placed at precisely defined locations. We further demonstrate that a bioink containing nonmotile bacteria can be used to create living boundaries that cannot be crossed by chemotactic bacteria, and thus can be used to guide chemotaxis within a hydrogel.

## 2. Results and Discussion

### 2.1 Encapsulation of Living *E*.*coli* Bacteria in Printable Hydrogel Matrices

In order to establish a reliable biofabrication procedure that enables reproducible printing of living bacteria into any desired target structure, several challenges have to be addressed: First, the printing process is required to induce only minimal stress on the bacteria, which favors printing procedures that work at low temperatures, low pressure, and do not involve radical crosslinkers. Second, a non-toxic hydrogel composition has to be found that has good printability with fast solidification and high structural fidelity. Lastly, after extrusion the printed bacterial gels have to be incubated for recovery and growth of the bacteria. To this end, the printed structures need to be embedded in a supporting environment that prevents shrinking, drying, or depletion of nutrients.

We first sought to find a printable hydrogel composition compatible with a moderate-temperature printing process using a custom-built gel extrusion printer (Figure 1A & Experimental Section),^[12]^ in which the highest continuously applied temperature does not exceed ≈ 42°C, and which allows for fast post-print solidification by cooling to room temperature (i.e., not requiring a crosslinking step). *E*.*coli* bacteria are known to display strongly reduced cell growth after exposure to elevated temperatures during growth^[13]^, with only some strains sustaining temperatures of up to 48.5°C ^[14]^. Higher temperature generally results in bacterial cell death or – in the case of spore-forming bacteria such as *B*.*subtilis*^[2a]^ – sporulation, which precludes a direct extrusion of living cells at elevated temperatures.

**Figure 1.**
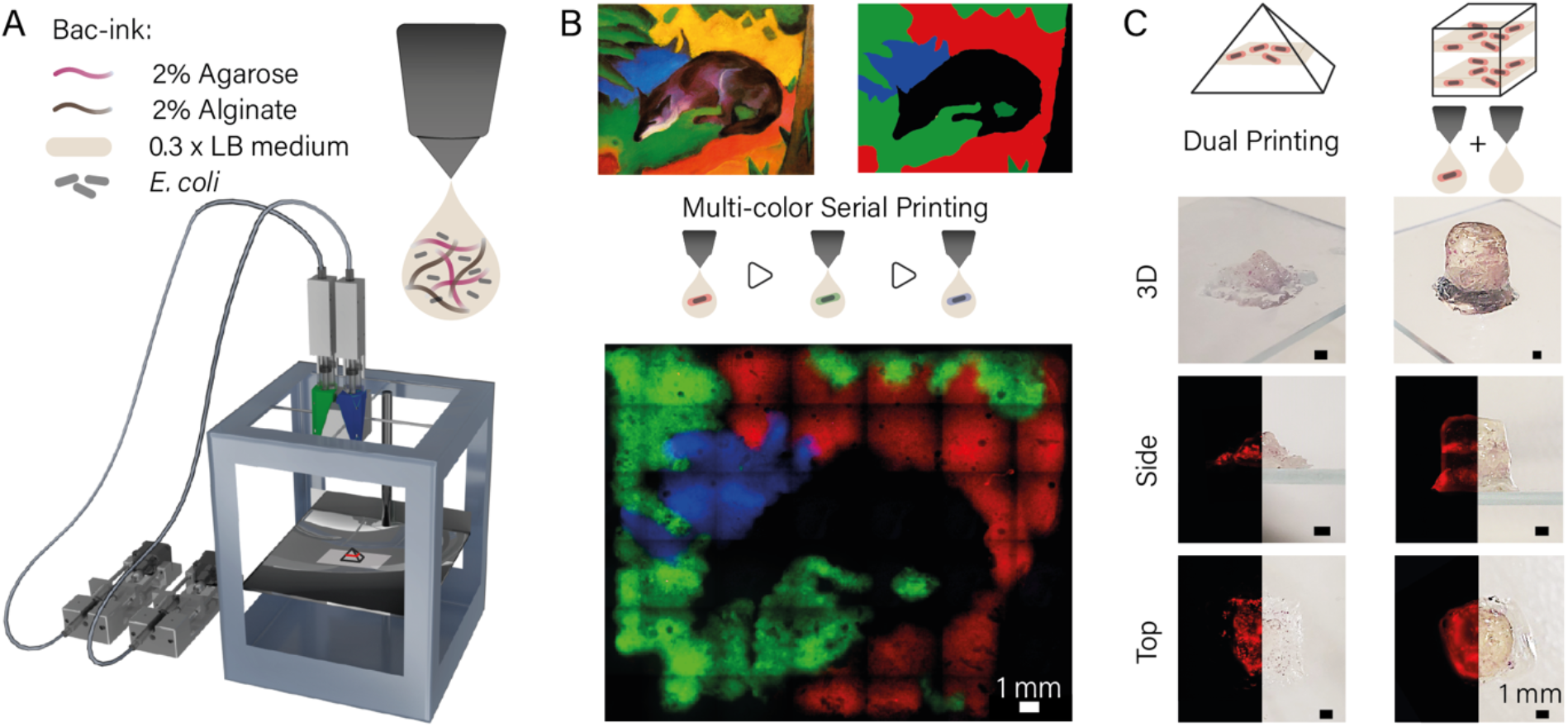
(A) Schematic image of a custom-built dual extrusion gel printer^[12]^ used for bacteria printing. Two syringe pumps were hydraulically connected to a print head holding two syringes, which were filled with bacteria-ink. The print heads were heated to 42°C, and the gel quickly solidified upon cooling on the base plate. (B) Bacteria (*E*.*coli* DH5α) expressing either mRFP, mVenus, or mTurquoise were printed sequentially into hydrogel layers and incubated over night to represent the painting “Blue Fox” by the Munich expressionist Franz Marc. The image demonstrates the high precision of bioink deposition and the reproducibility of printing at specific locations. C) Using dual extrusion with blank and bacteria-ink, three-dimensional gel objects were printed that contained single layers of bacteria. Images of the 3D structures (top and side view) were taken using a Canon 80D camera, fluorescence images were obtained with a Nikon Ti2-E microscope from the bottom and from slices of the structures.

We found a mixture of 2% agarose and 2% alginate to meet our requirements for a bacteria-ink, which are both hydrogels proven to be compatible with cell culture conditions ^[15]^. While alginate works as a viscosity enhancer, agarose provides fast solidification after extrusion and a good spatial resolution. To reduce osmotic stress, the bacteria-ink was prepared with 0.3 × LB medium which was found to be an LB concentration compatible with printing at 42°C.

Viscosity measurements indicate that the LB content has an appreciable influence on the temperature-dependent viscosity of the bacteria-ink mixture, which shows a steep decline in the temperature range from 20-35°C (SI Figure S1 and S2). During printing, the bulk of the gel is enclosed in the printhead, which works as a heat block at 42°C and fixes the temperature of the bacteria-ink to 42 ± 1.5°C within the sample syringe. By contrast, the tip of the nozzle is continuously exposed to room temperature, cooling the gel in this area to 36 ± 1.5°C during the printing procedure. The temperature of the extruded bioink droplets thus exactly matches the gelation temperature, while the main *E. coli* gel at 42°C provides tolerable conditions as well as a comparatively low viscosity, which improves printability and reduces the strain on the cells. At higher LB concentrations alginate crosslinking by the residual calcium in the medium results in gels with a larger variability in viscosity at the printing temperature, and lower viscosities at room temperature (SI Figure S1 and S2). With bacteria-ink containing 0.3 × LB medium, however, we were able to print structures up to 30 mm in size in x-y and 10 mm in the z dimension. Printing of larger structures was only limited by longer printing times, which resulted in drying of parts of the hydrogel during the printing process. As explained in more detail below, expression and maturation of fluorescent proteins inside the printed gel was supported by an oxygen-generating system composed of calcium peroxide and catalase.

### 2.2 Bioprinting of Living Bacteria in 2D and 3D

We assessed the printing performance of the bacteria-ink using various test structures. We first investigated the spatial position of the bacteria over time by imaging a printed triangular structure (Figure S3) with 4x, 10x and 60x magnification at a single position directly after printing and after 24h of incubation. The total fluorescence signal increased considerably due to the growth of the mRFP-expressing bacteria. Printed bacteria appeared to remain at or close to their initial positions over time (Figure S3).

In order to demonstrate the spatial resolution and reproducibility of the printing procedure in 2D, bacteria that constitutively expressed mRFP, mVenus and mTurquoise, respectively, were printed sequentially in the same plane (Figure 1B). After incubation at 37°C for 24h the gel image is clearly visible in the three fluorescent channels.

Overlapping signals from two fluorescence channels were found only in the border regions between subsequent print layers, which had widths between 50-500 µm, indicating the accuracy of the alignment of the printing steps with respect to each other.

We further used the dual extrusion capability of our printer to generate a pyramid with one and a cuboid with two distinct bacterial bands within a single print job (Figure 1C). Ink containing mRFP producing bacteria and bio-ink without bacteria were prepared and loaded into two separate glass syringes. The gels were printed alternatingly from the two separate syringes to form the 3D shapes with a base area of 6 mm × 6 mm each and a height of 5mm (pyramid) and 10mm (cuboid), respectively. Each bacterial band consisted of two layers printed with bacteria containing gel. After 24h of incubation at 37°C in LB medium in a 10% Pluronic F127 sealed chamber the printed structures were released from the Pluronic by dissolving the gel after 15 min storage at 4°C. The bacterial bands show high RFP production and are clearly colored red. Photography and microscopy images from the bottom and side of the prints demonstrate the precise deposition of the bacterial layers within the complete gel structure.

### 2.3 Growth Conditions in the Hydrogel Matrix

In order to gain a better understanding of the bacterial growth conditions inside the printed gels, we studied size and shape of the printed colonies for the first 8h after extrusion using confocal microscopy (Figure 2). The colonies typically exhibited ellipsoidal, in some cases spherical shapes, whose volume grew exponentially over time (Figure 2A). We converted the volume of the colonies into a cell count assuming a single cell volume of 1.5 µm^3^, which we also experimentally confirmed directly after the printing process. An exponential fit to the cell number as a function of time resulted in a mean generation time of 55 ± 6 min (cf. Figure S4 for experimental replicates).

**Figure 2.**
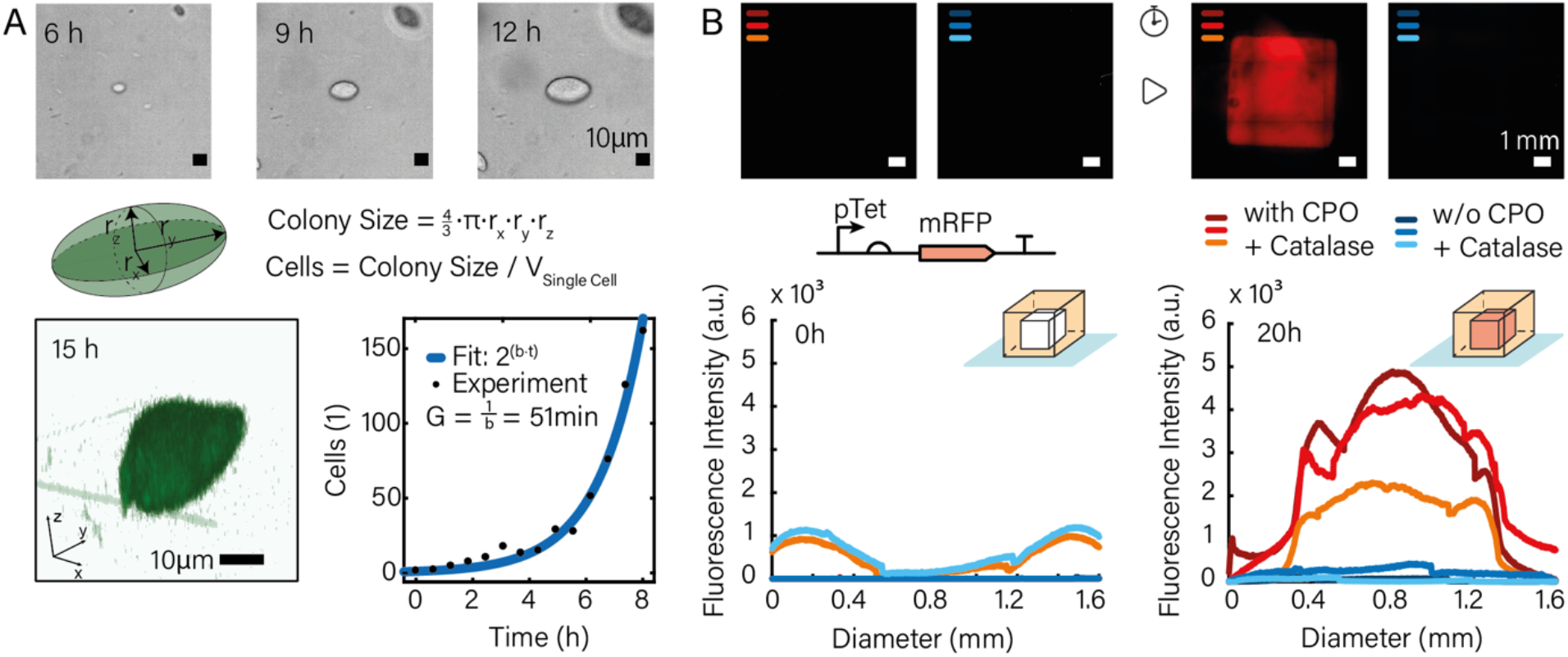
A) Bacterial growth observed by confocal microscopy. Bacterial colonies were generally found to be of ellipsoidal shape, whose sizes were used to determine bacterial growth curves inside the printed hydrogel matrix. The bacteria displayed exponential growth over at least 8 hours. B) For a good fluorescence readout, a continuous oxygen supply in the gel is necessary as maturation of the majority of fluorescent proteins depends on oxygen ^[6, 16]^. We casted two cuboid structures and sealed both in LB-ink to mimic the procedure applied for printed structures. With the addition of 0.15% calcium peroxide and 0.3 mg/ml of the enzyme catalase the fluorescence signal strongly increased during 20h of growth at 37°C while in the absence of any oxygen generator, no fluorescence could be detected.

In previous work on bacterial bioprinting ^[2a]^, expression of fluorescent proteins was only observed at gel-air interfaces, which is explained by the lack of oxygen – required for FP maturation – inside the gel. As our studies required fluorescence monitoring also of bacteria deeply embedded in the gel, we utilized a calcium peroxide-based oxygen generation system, which is known as an additive from conventional cell culture ^[10]^. Within the gel, CPO degrades to hydrogen peroxide, whose conversion to oxygen and water is then catalyzed by the enzyme catalase. As shown in Figure 2B, we casted (6 mm)^3^-sized hydrogel cubes with bacteria containing the plasmid pTet-mRFP and induced the expression of mRFP via addition of anhydrotetracycline (aTc) directly before mixing with bacteria-ink either with or without the addition of CPO and catalase. While after 20 h of incubation the sample without CPO did not provide an appreciable fluorescence signal, the gels containing the oxygen generating system showed a clear fluorescence signal in all triplicates.

### 2.4 Diffusion of Genetic Inducers

We next studied the diffusion of inducer molecules and the resulting spatiotemporal gene expression response of bacteria embedded in the bioprinted gel, for which we conducted experiments with aTc-inducible DH5αZ1 cells carrying a plasmid that encodes mRFP under the control of a pTet promoter (Figure 3A and B). The cells were printed into a circular shape with a diameter of 12 mm. Directly after printing, a 2 µl voxel containing aTc (final concentration in the reaction chamber 0.2 µg / ml (or 0.43 µM)) was added to the center of the structure directly after the print. The printed structure was then sealed and imaged continuously for 12h at 37°C. After incubation for 3 – 4 hours, the fluorescence signal started to rise in the center while small colonies became visible in the brightfield images. After incubation for 10 hours, large bacterial colonies were visible in the brightfield channel and the fluorescent protein was highly expressed by the bacteria.

**Figure 3.**
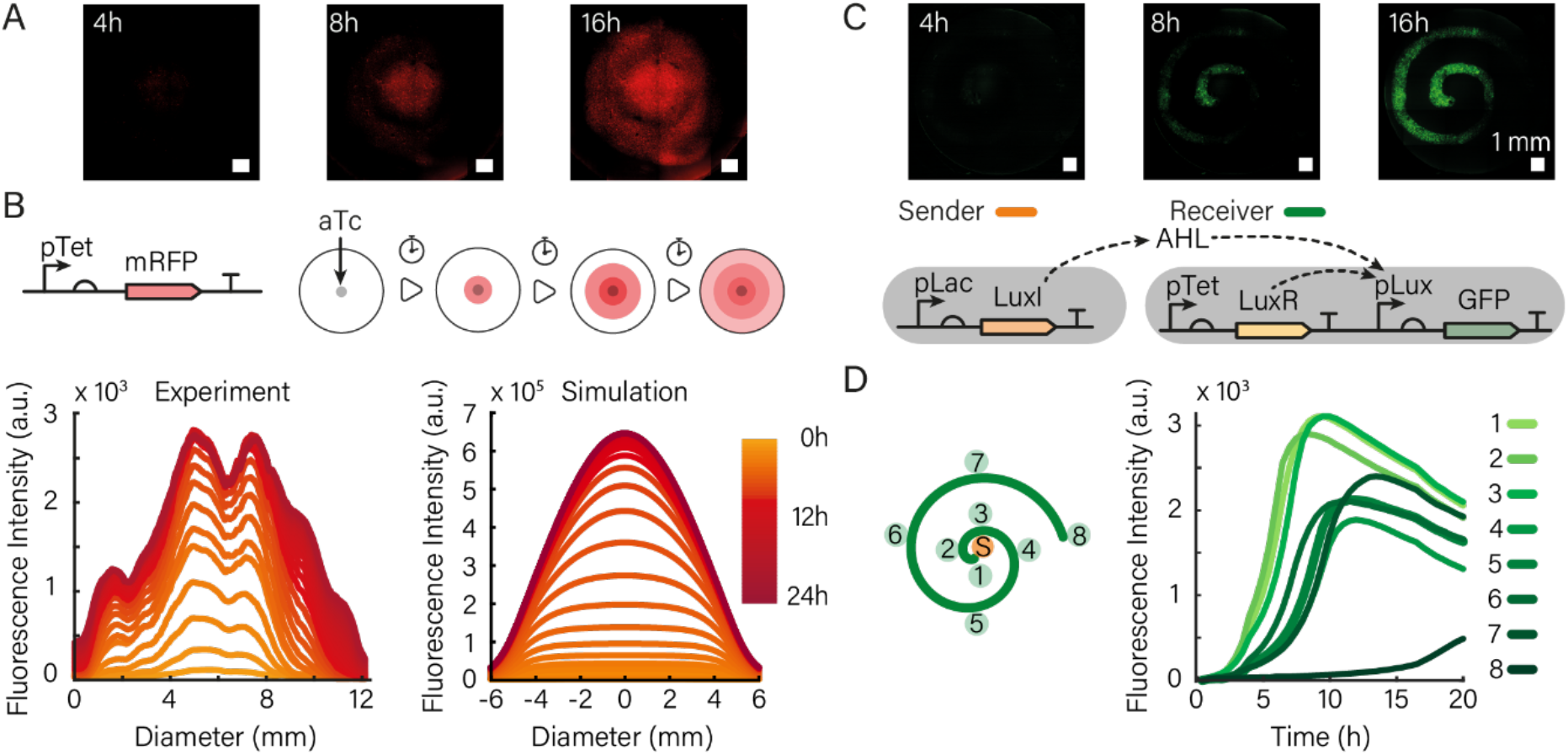
A) Diffusion of the genetic inducer aTc from the center of a printed gel and subsequent expression of mRFP by bacteria contained in the gel (scale bar 1 mm). B) mRFP was expressed under the control of a pTet promoter and thus responds to aTc diffusion. The graph on the left shows the fluorescence intensity measured along the diameter of the bioprint for different timepoints. Simulations (right) that model aTc diffusion, bacterial growth and gene expression are in good agreement with the experimental observations. C) Fluorescence channel of 4x microscopy images of a sender-receiver system at 4h, 8h, and 16h after the start of the experiment. The increase in fluorescence intensity is synchronous (time intervals of 10 min showed no differences) upon induction of the sender bacteria with IPTG and consequently with high AHL production. The corresponding gene circuit is shown below the micrographs. D) Fluorescence time traces (right) corresponding to the locations indicated on the left for an experiment, in which the sender bacteria were not induced, and thus only a small amount of AHL is produced by leaky expression of LuxI from the Lac promoter.

The observed spatiotemporal dynamics of the fluorescence signal in the gel result from a combination of the diffusion of the inducer from the center into the gel print, bacterial cell growth throughout the print, and inducer concentration-dependent gene expression in the bacteria. As shown in Figure 3B the rise of the fluorescence signal can be well recapitulated in a numerical simulation of these processes using experimentally realistic parameters for bacterial growth rate and expression dynamics (cf. Figure S4 - S6, and Supporting Information Section 4.1 on the numerical model). Screening of the value of the diffusion coefficient for aTc in simulations suggest a diffusivity on the order of 200 µm^2^/s within the gel (SI Figure S10).

### 2.5 Bacterial Communication in a Bioprinted Environment

Bioprinting with bacterial inks opens up the possibility to arrange different types of communicating or otherwise interacting bacteria into well-defined spatial relations. This is of particular interest for applications in synthetic biology, which require the precise control of the boundary conditions affecting the execution of spatiotemporal gene circuits^[11a-d]^ by engineered bacteria. As an example for this type of application, we studied the dynamics of a synthetic bacterial sender-receiver system embedded in a bioprinted gel structure. The sender cells are DH5α cells hosting a plasmid coding for the 3OC6-HSL (AHL) synthase LuxI under the control of a Lac promoter (cf. Materials & Methods). Upon induction with IPTG the cells express LuxI which consecutively produces the quorum sensing signal AHL that can freely diffuse out of the bacteria and through the gel. The corresponding receiver cells contain a plasmid for the constitutive production of the AHL-dependent transcriptional activator LuxR, which controls expression of GFP via the pLux promoter. The basic function of the sender-receiver system was first checked in bulk experiments using a plate reader (SI Figure S7).

In order to illustrate spatial control over sender-receiver dynamics, we then printed the receiver cells in the form of an Archimedean spiral (defined by r(ϕ)= a ϕ, where r, and ϕ are polar coordinates, and a is a constant) and positioned the sender cells at its center (r=0).

As expected, AHL produced by the senders activates production of GFP by the receivers along the spiral (Figure 3C, Supporting Video 1). The signal typically increased over 10-16 h until nutrient depletion set in, followed by a decay in GFP fluorescence.

The behavior of the sender-receiver system differs from that of the aTc-inducer system due to the much higher sensitivity of LuxR activation to AHL (with a K_d_ on the order of only 10 nM). We therefore performed experiments both with fully induced senders, but also with uninduced senders that generated AHL via leaky expression of LuxI (and thus had a correspondingly lower “sender strength” ^[17]^).

We found considerable variability among the sender-receiver experiments (SI Figure S8), which is likely related to the sensitivity of the dynamics with respect to variations in bacterial growth and the very low AHL concentration required for induction (cf. discussion in Section 4.2 of the Supporting Information). At low sender strength, in some cases there was a clear differentiation in spatial response of the bacteria, with the closer receiver bacteria responding faster than the more remote ones (Figure 3D). In other cases, there was no clear spatial order, or the complete receiver spiral started to produce GFP synchronously (Figure S8). Simulations indicate that in this case the AHL concentration throughout the gel had already reached nanomolar concentrations before the receivers started to grow appreciably (SI Figure S11).

### 2.6 Guiding Chemotaxis with Printed Boundaries

As a second example for spatial control over bacterial dynamics provided by the bacterial bioprinter, we investigated the motion of chemotactic bacteria in engineered environments. Chemotactic bacteria can move on top of (and to a certain extent also underneath) the surface of soft agar gels ^[18]^, propelling themselves towards areas of more favorable environmental conditions. For our experiments, we prepared soft agar plates at agar concentrations between 0.22-0.24% (w/v) into which we printed artificial boundaries in the shape of our university logo formed by RFP producing non-motile *E*.*coli* cells mixed with 1% alginate as printable ink (Figure 4A). The plates were then inoculated with 2µl aliquots of a suspension of a chemotactic *E. coli* strain (MG1655) placed at approximately equal distances from the outline of the logo. Within 20h the chemotactic cells grew into a thick biofilm over the agar surface, avoiding only the area enclosed by and between the printed borders.

**Figure 4.**
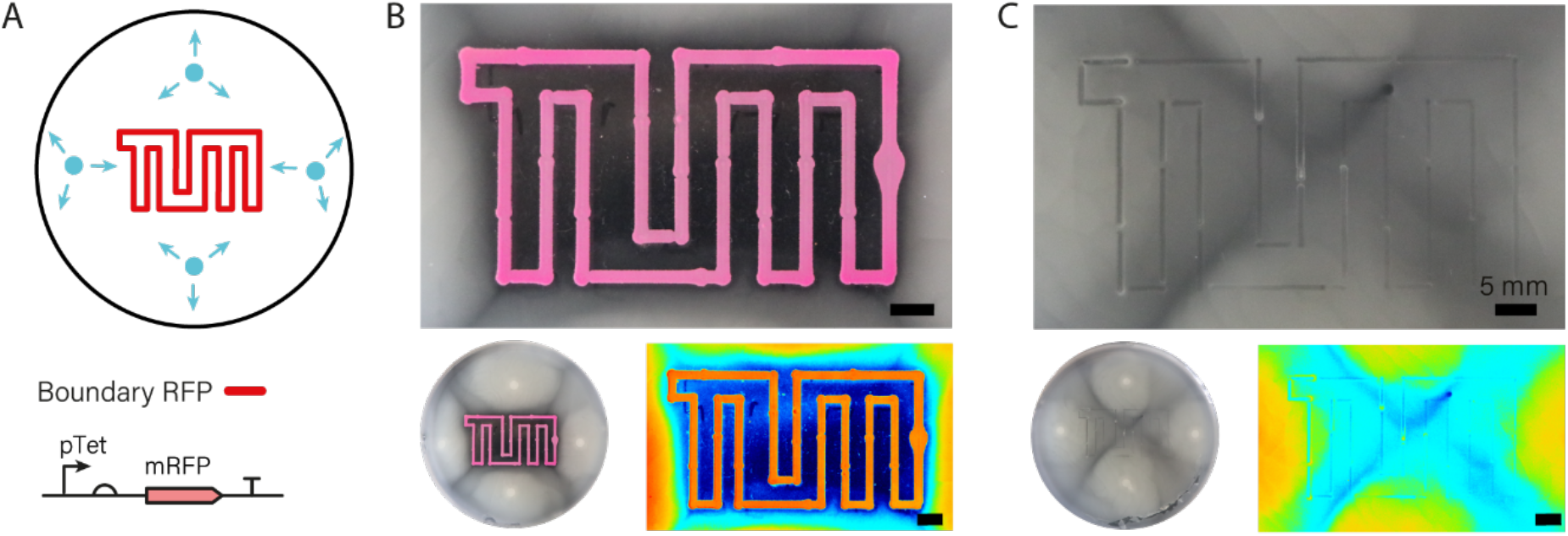
A) Artificial boundaries in the form of a TUM logo were printed into soft agar using non-motile bacterial cells constitutively expressing RFP. Directly after printing, chemotactic cells were inoculated on four locations and grown at 37°C. B) After 20h of incubation, the printed boundaries showed a high intensity of RFP fluorescence. The chemotactic cells successfully spread over the whole surface outside of the boundaries, but could not enter the interior of the logo. The small images show a view of the whole plate (left) and a high contrast image of the apparent bacterial density in heatmap coloring (right). C) For comparison, the TUM logo was printed with bioink not containing any bacteria. In this case the bacteria spread over the whole plate without hindrance.

Apparently, the presence of the printed bacterial border region prevented the chemotactic bacteria from propagating into the interior of the logo, which was found to be only scarcely populated (Figure 4B). We surmise that the chemotactic bacteria do not enter the border region due to the unfavorable nutrient conditions generated by the growing border bacteria, potentially combined with the physical obstruction caused by the non-motile bacteria themselves. In control experiments, in which a bacteria-free alginate boundary was printed, the chemotactic bacteria were able to completely fill the whole agar plate (Figure 4C).

## 3. Conclusion

In the present work we have demonstrated a viable approach towards 3D printing of living hydrogel materials using bacteria-laden bioink. The mild low-temperature extrusion conditions of our protocol combined with a non-toxic hydrogel ensured survival of the bacteria throughout the printing process and thereafter. *E. coli* cells were shown to exponentially grow into extended microcolonies within the gel. Addition of an oxygen generating system based on calcium peroxide and catalase also enabled the expression of fluorescent proteins deep within the gel. We showed the possibility of dual extrusion of hydrogels into 3D shapes and sequential extrusion of hydrogels containing different types of bacteria, both with a high spatial resolution. Notably, the ability to print living *E*.*coli* bacteria opens up the possibility to spatially organize genetically engineered, communicating and interacting bacteria, which is of great interest for applications in synthetic biology. We demonstrated this capability by showing spatiotemporal gene expression dynamics controlled by diffusing inducer molecules, or generated by a bacterial sender-receiver system. We also showed that bacteria ink loaded with non-motile bacteria can be used to define and grow living boundaries, which are impenetrable for motile bacteria co-embedded in the hydrogel.

We anticipate that the spatial organization of live bacteria by 3D bioprinting will enable the generation of soft living biomaterials that are capable of simple forms of differentiation. In such materials, bacteria can sense environmental inputs and produce molecules in response to these molecules in a spatially differentiated manner. Communication between bacteria further supports spatial organization and division of labor between the bacteria. Extrusion of different types of bacteria could even be used for the spatial organization of co-cultures which cooperatively perform more complex biosynthetic tasks.

## 4. Experimental Section/Methods

### Bioprinter

Bioprinting of bacteria-containing hydrogels was performed using a custom 3D printer setup based on a commercially available Ultimaker Original+ modified with custom-built components ^[12]^. Hydrogel is extruded from heated printheads via hydraulically coupled syringe pumps. In order to increase the variety of structures that can be manufactured, the printer was equipped with two extruders. The extruders can be loaded with any pair of hydrogels that can be printed at the same temperature. Both extruders are directly controlled by the Ultimaker board. Executable GCODEs for all structures were generated by our Python based software (available on https://github.com/julia-mueller/bioprinter/).

### Bacterial strains and culture conditions

For all experiments *E*.*coli* strains (DH5α, DH5αZ1, BL21, MG1655) with different plasmids were cultured from glycerol stocks and grown over night in LB medium containing selective antibiotic (carbenicillin, Carb) in an incubator shaker (Innova44, New Brunswick) at 37°C and 250 rpm. On the following day, cells were diluted 1:100 in LB medium containing carbenicillin and cultivated to an OD of 0.6. Prior to printing, the cells were centrifuged at 2000 rcf at 4°C for 2 min and the pellet was resuspended in fresh LB medium containing carbenicillin.

The chemotactic bacteria used in Fig. 4 were MG1655, while the non-motile bacteria were DH5α.

### Plasmids coding for fluorescent proteins

High copy number plasmids (iGEM part pSB1A3) containing a constitutive promoter (iGEM part J23106) and the coding sequence for mVenus (YFP), or mTurquoise-2 (CFP), were transformed to DH5α cells for constitutive protein expression. The coding sequence for mRFP under a pTet promoter (iGEM part I13521) on a high copy number plasmid (iGEM part pSB1A3) was transformed to DH5alpha, which does not express a TetR, for ‘constitutive’ and to DH5αZ1 cells for aTc inducible protein expression.

### Sender-receiver plasmids

The sender plasmid (iGEM part K116638) is a high copy number plasmid (iGEM part pSB1A2) and contains a pLac promoter (iGEM part R0010) controlling the AHL synthase LuxI (iGEM part C0061) and was transformed into DH5α cells for IPTG inducible AHL expression. Leaky expression of LuxI and thus production of AHL can occur as there are small amounts of Lactose present in LB medium. The receiver plasmid is a high copy number plasmid (iGEM part pSB1A3) that contains an expression cassette coding for the transcription factor LuxR (C0062) regulated by a Tet promoter (iGEM part R0040), and a second cassette coding for GFP (iGEM part E0040) regulated by a AHL inducible Lux promoter (pLux). We transformed this plasmid to BL21(DE3)pLysS.

### Bacteria-ink

For our bacteria-ink we used final concentrations of 2% agarose and 2% alginate in 0.3 × LB medium containing (final) concentrations of 0.5% calcium peroxide (CPO) and 300µg/ml catalase.

### Bioprinting of bacteria

Mixed bacteria-ink was prepared in a microwave oven and kept at 70°C for 15 min in an oven prior to mixing with bacteria culture. 500µl of a liquid bacterial culture at OD 0.6 was mixed with 1 ml of bacteria-ink that was enriched with 60µl catalase (end concentration 300µg/ml) and 200µl CPO (end concentration 0.05% w|v). The mixed bacteria-ink was loaded into the printhead that was preheated to the printing temperature. Extrusion on glass slides (microscopy slides 76mm × 26mm × 1.5mm, VWR, Germany, and Menzel slides 76mm × 26mm #1.5 Thermo Scientific, German) was performed with nozzle No. G23 (ID 330 μm) (Vieweg, Germany). For dual extrusion, a second gel was prepared and loaded into a second glass syringe in the printhead. Prints on agar plates were performed with a mixture of 500 µl of liquid bacterial culture at OD 0.6 and 500 µl 2% (w|v) alginate loaded to the printhead and extruded at 37°C with nozzle No. G25 (ID 250 µm) (Vieweg, Germany).

After printing, the extruded structures were sealed with a custom 3D printed plastic frame, LB-bacteria-ink and closed with a glass lid. Directly after sealing, imaging with an epifluorescence microscope (Nikon Ti2-E) was started. For different magnifications a 4x, 10x, and 60x objective were employed. For larger structures (>1 mm), multipoint imaging with 4x magnification and consecutive stitching with 0% overlap of single images was performed.

### Sample sealing

All samples contain between 0.1 ml and 0.3 ml bacteria-ink extruded in thin lines and are thus extremely prone to drying and shrinking during incubation at 37°C. As drying of the hydrogel leads to cell death, drying and consecutive shrinking was prevented using custom 3D printed plastic chambers (polylactic acid, BASF, Germany) and glass cover slides (round 18mm diameter and square 15mm, #1.5, Carl Roth, Germany) that are attached to the cover slides with instant glue.

### Large sample stabilization

For stabilization during incubation of larger structures (pyramid and cuboid with bacterial layers) the chambers were filled up with 10% Pluronic F127 (Sigma Aldrich, Germany) in LB medium. This inverse thermostable polymer stabilizes higher structures as its viscosity enhances extremely while incubating at 37°C and can be removed as it liquefies after 15min at 4°C. As shown in early work on bacteria cultures on Pluronic F127 as agar substitute, bacteria are able to grow on this polymer ^[19]^. However, higher concentrations of Pluronic lead to cell death if the cells are exposed to Pluronic for several days (this polymer is also used as an anti-fouling agent used for long-term anti-biofilm surface treatment ^[20]^). With a maximum of 24h incubation of bacteria-ink structures using low concentrations of this stabilizing agent, we ensured cell growth on the printed hydrogel structure without contamination of the surrounding auxiliary structure by washed out cells ^[21]^.

### Nutrient and oxygen supply

To prevent drying and to ensure a continuous supply with nutrients we embedded all printed structures in bacteria-ink in 1x LB medium which provides enough nutrients to feed the enclosed bacteria for at least 24 hours. Sealing of the structures reduces the supply of the samples with oxygen, which is required, however, for the maturation of the fluorescent proteins used for optical readout ^[16b, 22]^, in particular for red fluorescent proteins ^[16a]^. We therefore augmented our bacteria containing bioink with 0.05% (w/v) calcium peroxide (CPO) and 300 µg/ml catalase, which has been found sufficient to supply cell cultures with oxygen for over a week ^[10c, 23]^. CPO slowly decays into hydrogen peroxide at room temperature while catalase promotes the conversion of H_2_O_2_ to oxygen and water.

## Supporting Information

Supporting Information is given at the end of this manuscript

## Acknowledgements

The authors thank B. Buchmann from the group of Prof. A. Bausch for his help with rheology measurements and confocal imaging. This work was funded by the European Research Council (project AEDNA, grant no. 694410). We gratefully acknowledge funding by the Bavarian Ministry of Science and the Arts through the ONE MUNICH Project “Munich Multiscale Biofabrication”.

## 1. Experimental Protocols

### 1.1 Preparation of Bac-Ink

Composition of bac-ink:

50 ml ddH2O

1.5 g agarose NEEO quality (2267.3 Carl Roth, Germany)

1.5g alginate (G1890, Sigma Aldrich, Taufkirchen, Germany)

Protocol:

- heat ddH_2_O to 100°C
- add agarose while stirring at 1400 rpm for ≈ 10min
- add alginate slowly while still stirring (min 1400 rpm) for 45min
- degas at 80-100°C for 1 h

Bac-ink can be stored in closed bottles at room temperature and melted in a microwave prior to printing.

### 1.2 Bacterial Cell Culture

- 5ml LB medium with 5 µl carbenicillin (stock concentration: 1000x) and bacteria from glycerol stock
- Incubate for 16 h at 37 °C
- Dilute 1:100
- For printing set OD to 0.6 in fresh LB best: centrifuge suspension at 4 °C, for 2 min at 2000 rcf, resuspend with fresh LB media

### 1.3 Bioprinting

- Melt bacteria-ink in a microwave and keep warm in the oven at 70°C
- Preheat extruder and sample syringes at 42°C (printing temperature)
- Mix sample ink:
- 1 ml bacteria (OD 0.6)
- 2 ml bac-ink
- 60 µl catalase (stock:1 mg/ml ≙ 60 - 100 units/ml)
- 200 µl CaCO_2_ (stock: 0.15% w/v)
- Transfer 1 ml sample ink to preheated syringe
- Print on cover slide or microscopy slide (printhead: 42°C, printbed: 23°C, room temperature)
- Glue plastic frame (PLA, 3D printed) with inner cavity according to print dimensions (circular: diameter 17mm × 4 mm, rectangular: 24 mm × 46 mm × 4mm) around the print for sealing (instant glue)
- Chamber sealing: add 1ml to a circular or 6 ml LB-bacteria-ink to a rectangular chamber, glue #1 cover slide on top of the plastic chamber

Software and printer hardware are custom built and published already [5]. The complete code is available under a Creative Commons License on Github:

https://github.com/julia-mueller/bioprinter/

### 1.4 Epifluorescence Microscopy

Fluorescence intensities were captured for up to 24 h in single images every 10 – 30 min at 37°C with a P-Apo 4x NA 0.20, P-Apo 10x NA 0.45, or P-Apo 60x Oil NA 1.4 objective on a Nikon Ti-2E, equipped with a SOLA SM II LED light source, a motorized stage, perfect focus system, an Andor NEO 5.5 camera and the filter sets listed in Table T1. Typical settings were 50% brightness of the fluorescence LED and the exposure time given in Table T2. Imaging was performed at locations in close proximity with no overlap between neighboring images.

The captured images were then stitched together to cover the whole sample area. For inducer reaction-diffusion experiments 4 × 4 (Figure 4), for the sender-receiver experiment (Figure 4) 5 × 5, and for the artprint (Figure 2) 7 × 11 images were captured and stitched without overlap.

**Table S1.**
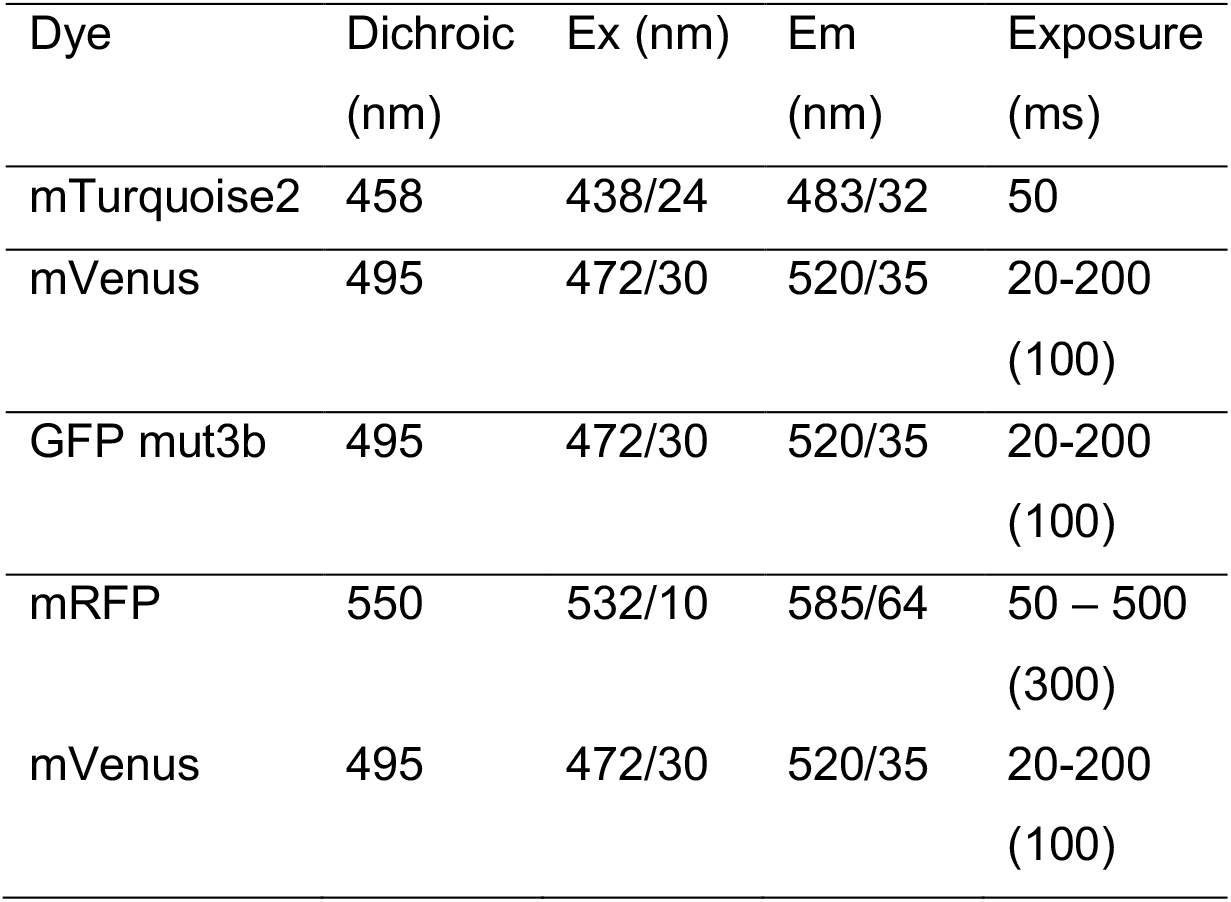
Filters and exposure times used in epifluorescence microscopy.

### 1.5 Confocal Microscopy

Fluorescence intensities were obtained from single images captured every 20 min and for up to 20 h at 37°C with an HCX PL APO CS 63.0x 1.40 UV oil objective on a Leica Sp5 confocal microscope equipped with an Ibidi incubation chamber. Samples were excited at 633 nm and fluorescence was collected with a photo-multiplier between 640 nm and 710 nm. Imaging was performed at specific locations with z-stacks with a height between 80 µm – 100 µm in 3.00µm steps.

### 1.6 Bulk Rheology

To measure the temperature dependence of the viscosity of the bioink, a TA-rheometer (MCR 301, Anton Paar) was used with a 50-mm plate-plate geometry. The sample volume was 750 µl for each measurement. All samples were heated to 70°C before transferring them onto the rheometer measuring plate. The rheometer was preheated to 60°C and each sample was measured for 1min at 60°C to ensure a constant temperature of 60°C for the whole sample volume. After this incubation time the measurement was started. The temperature was decreased in 1°C steps from 60°C to 20°C in 4s intervals.

## 2 Bacteria

**Table S2.** Sender - Receiver Systems.

| Name | Plasmid | Biobrick |
| --- | --- | --- |
| Sender | DH5α - pSB1A2 -<br>K116638 | iGEM part K116638 on parts.igem.org |
| Receiver | BL21pLys - psB1C3 -<br>T9002 | iGEM part T9002 on parts.igem.org |

**Table S3.** Fluorescent Bacteria

| Name | Plasmid | Biobrick |
| --- | --- | --- |
| mRFP | DH5α - pSB1A3 -<br>I13521 | iGEM part I13521 on parts.igem.org |
| mRFP | DH5αZ1 - pSB1A3 -<br>I13521 | iGEM part I13521 on parts.igem.org |

**Table S4.**
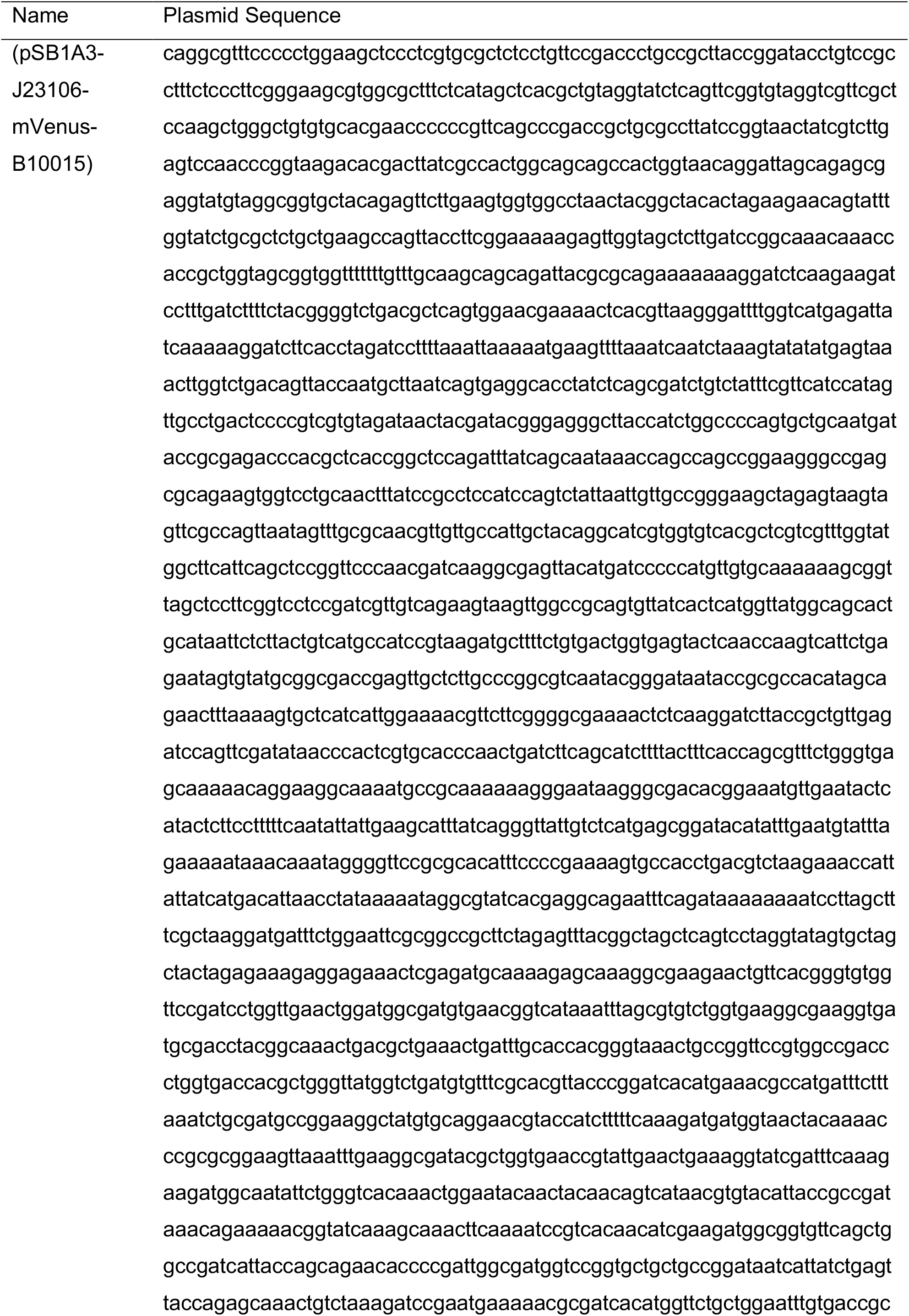
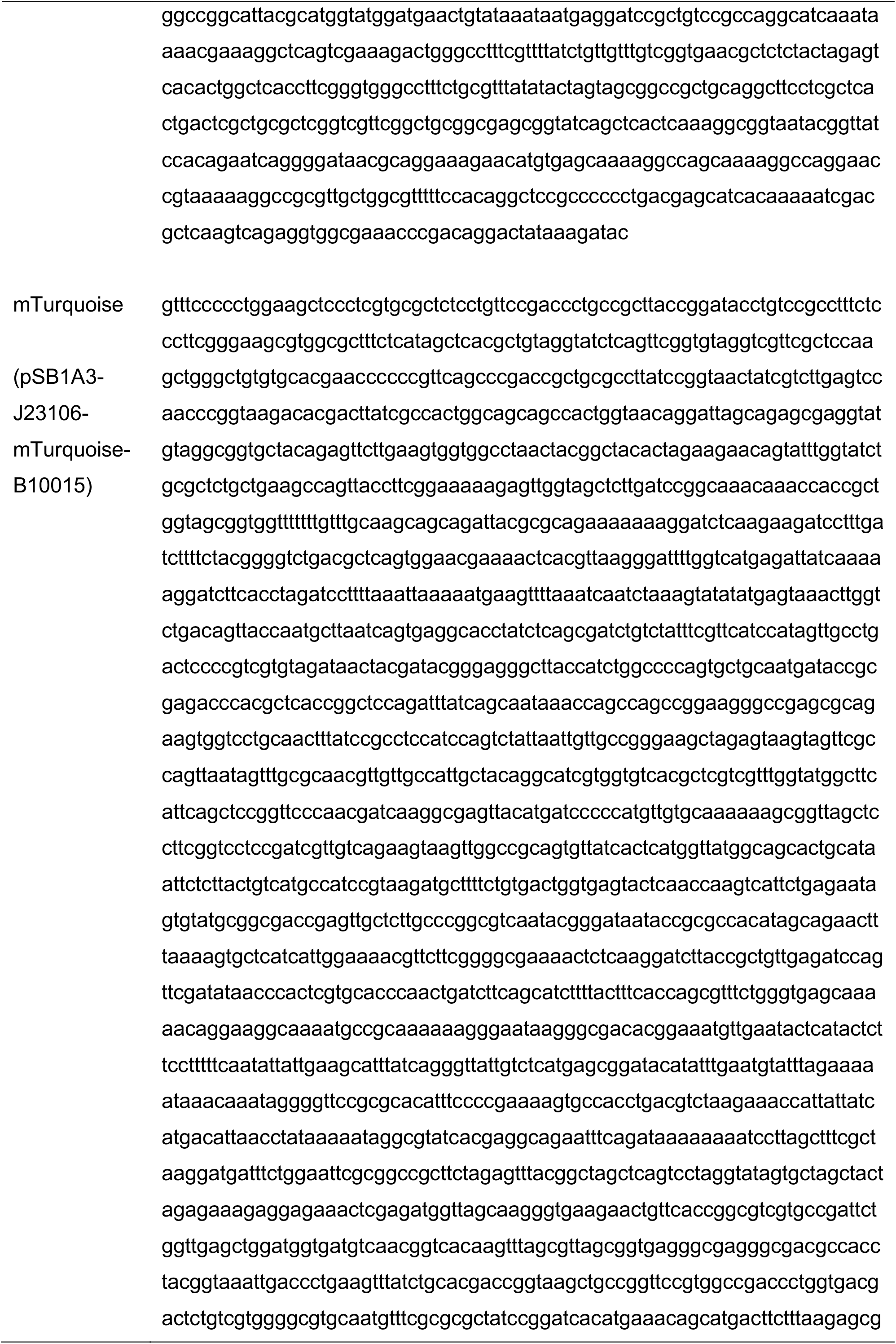
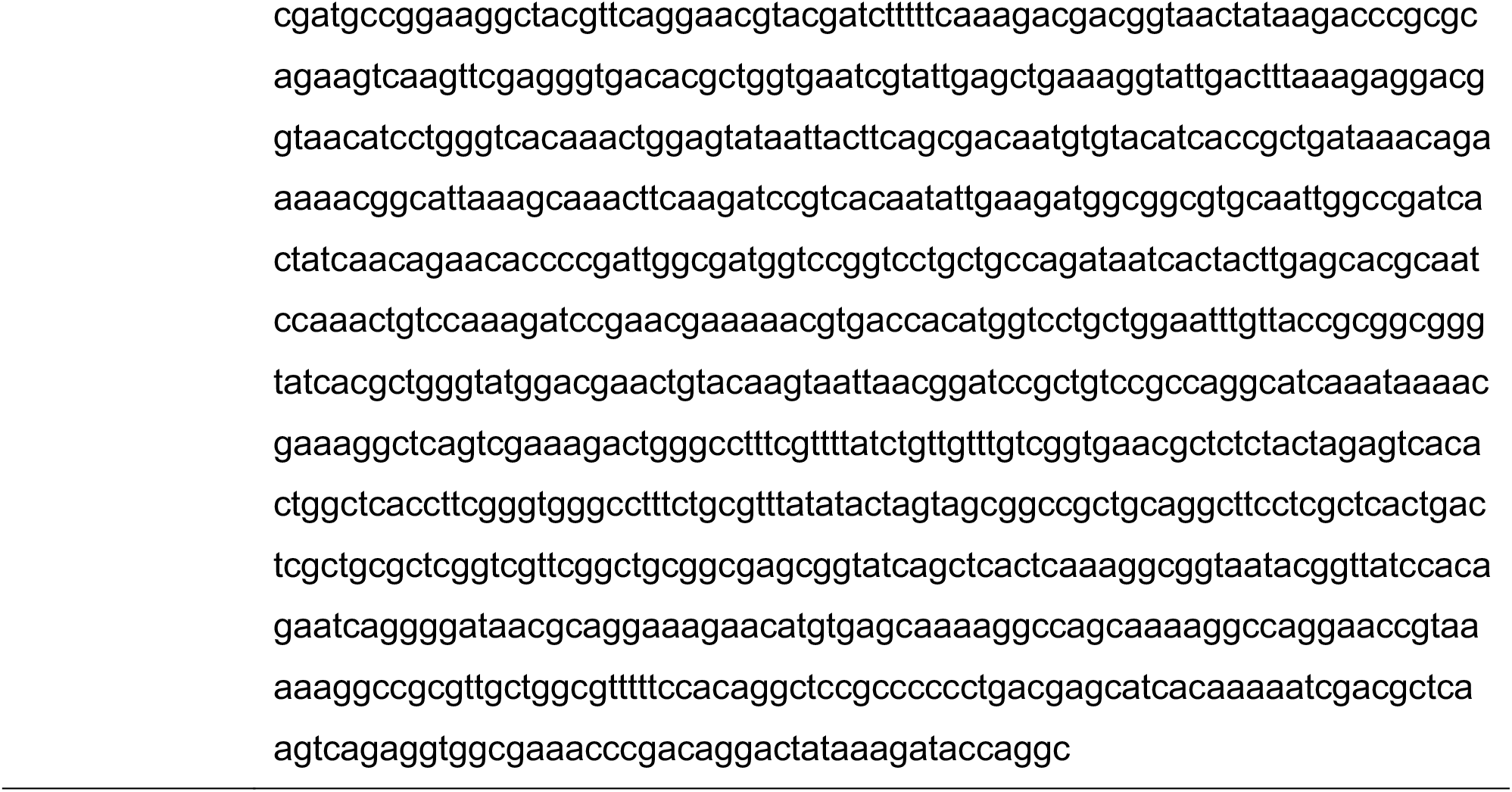
Fluorescent bacteria - plasmid sequences.

## 3. Additional Experimental Data

### 3.1 Bulk Rheometer Data

**Figure S1.**
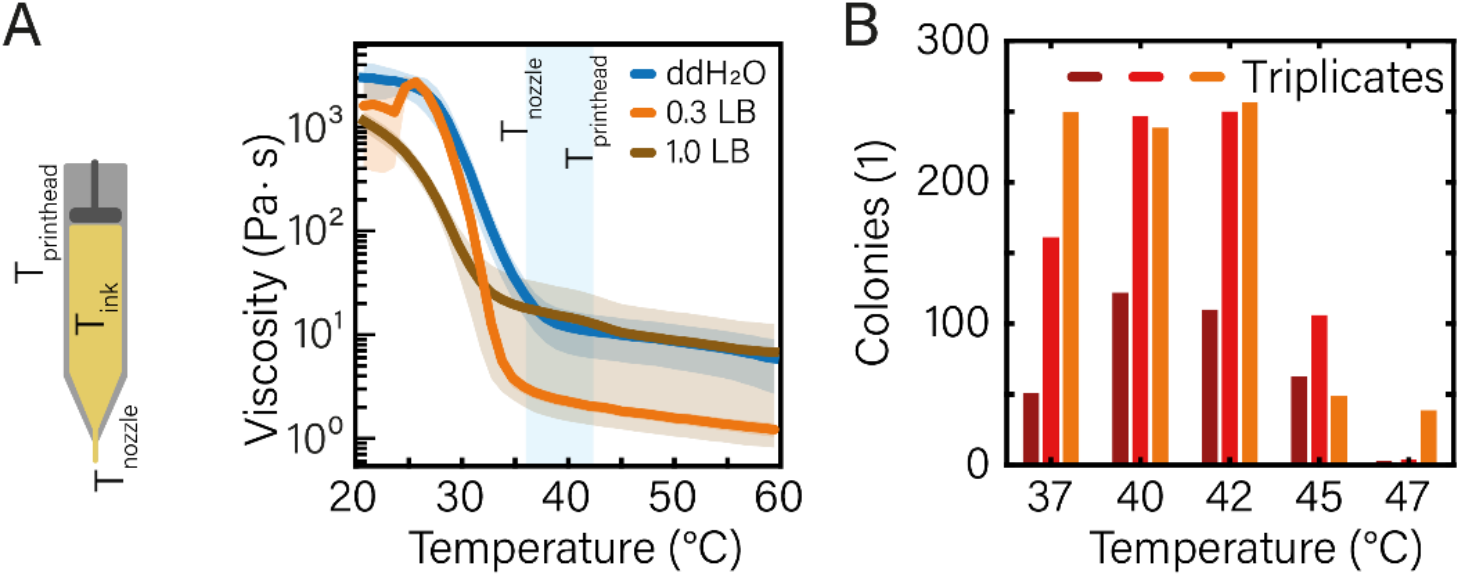
(A) Viscosity of bacteria ink prepared in either pure ddH2O, 0.3 × LB, or 1 × LB medium. We see that the region of interest is the temperature range between nozzle (≈ 36°C) and printhead (≈ 42°C) temperature. Fast solidification at all temperatures below the nozzle temperature is beneficial while solidification during the print leads to clogging. For bacteria ink in 1.0 × LB medium the viscosity is significantly lower at room temperature than for ddH_2_O or up to 0.3 × LB medium in ddH_2_O. This leads to low printability and less stability of extruded structures. (B) Bacterial growth after heat shock. The heat shock induced on the cells has a duration of maximum 2 min while the hot gel (60°C) is mixed with the liquid bacterial cultures (37°C) in a ratio 2:1. Afterwards the bacteria are kept at printing temperature for up to 30 min and can grow in the incubator at 37°C after printing. The preparation-induced heat shock is simulated with liquid bacterial cultures in a thermocycler. The cultures are plated after the heat shock and CFU were counted. Up to printing temperatures of 42°C the bacterial growth is similar or better than for the printing temperature (37°C). Printing temperatures higher than 47°C appear are not compatible with survival of the cells. These temperatures are consistent with earlier findings for heat stressed *E. coli* bacteria (not in the context of bioprinting).

**Figure S2:**
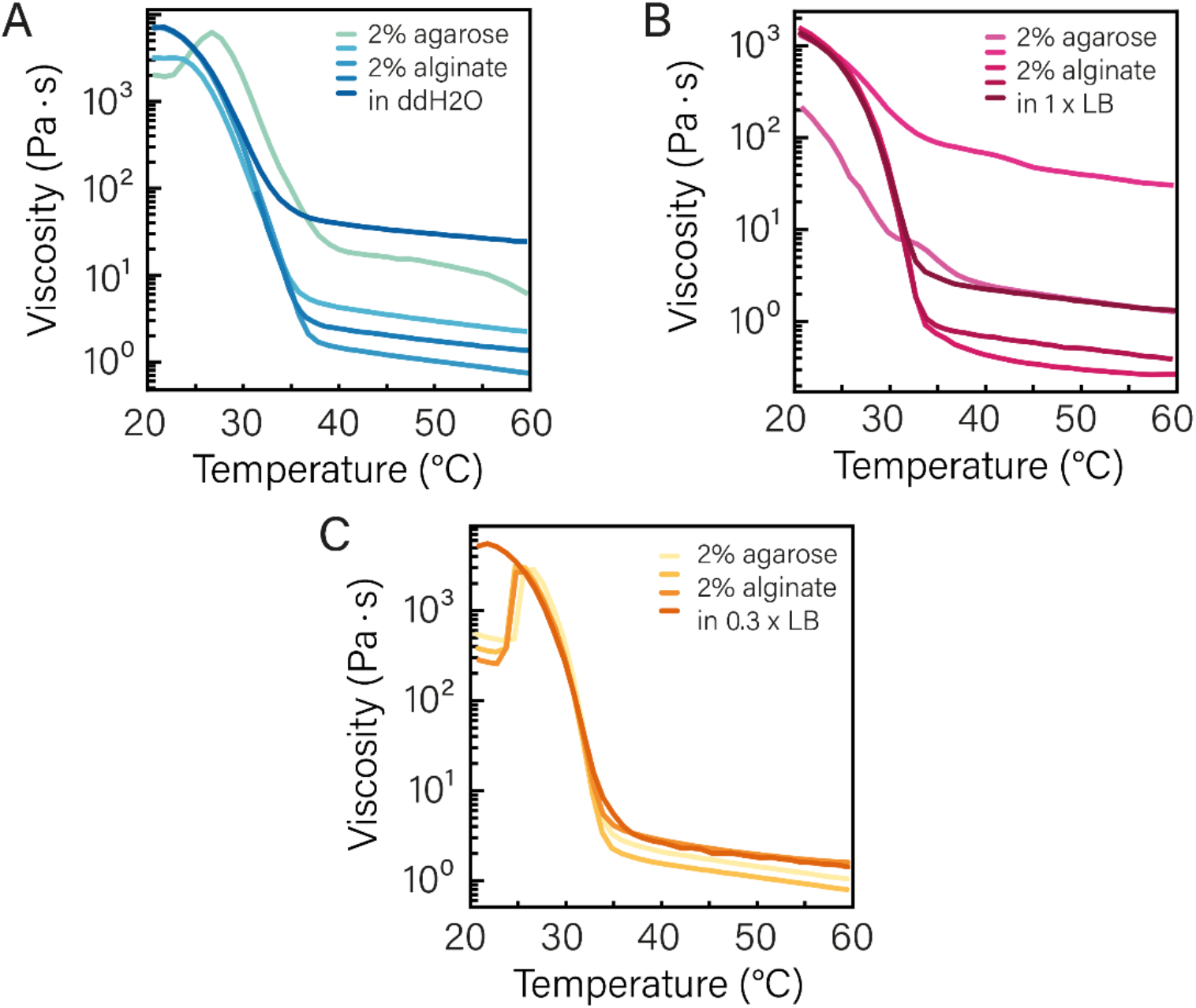
Experimental triplicate data for viscosity measurements on bac-ink in ddH_2_O, 0.3 × LB and 1x LB medium (cf. Fig. S1). The temperature dependence of the viscosity has a clear transition temperature for all conditions, which in all cases is below 35±2°C. Hence, the gels solidify at room temperature. For ddH_2_O and 1x LB medium the batch-to-batch variation is higher than for 0.3x LB medium, which also performed best in all prints.

### 3.2 Localization of Bacteria in Printed Gels

**Figure S3:**
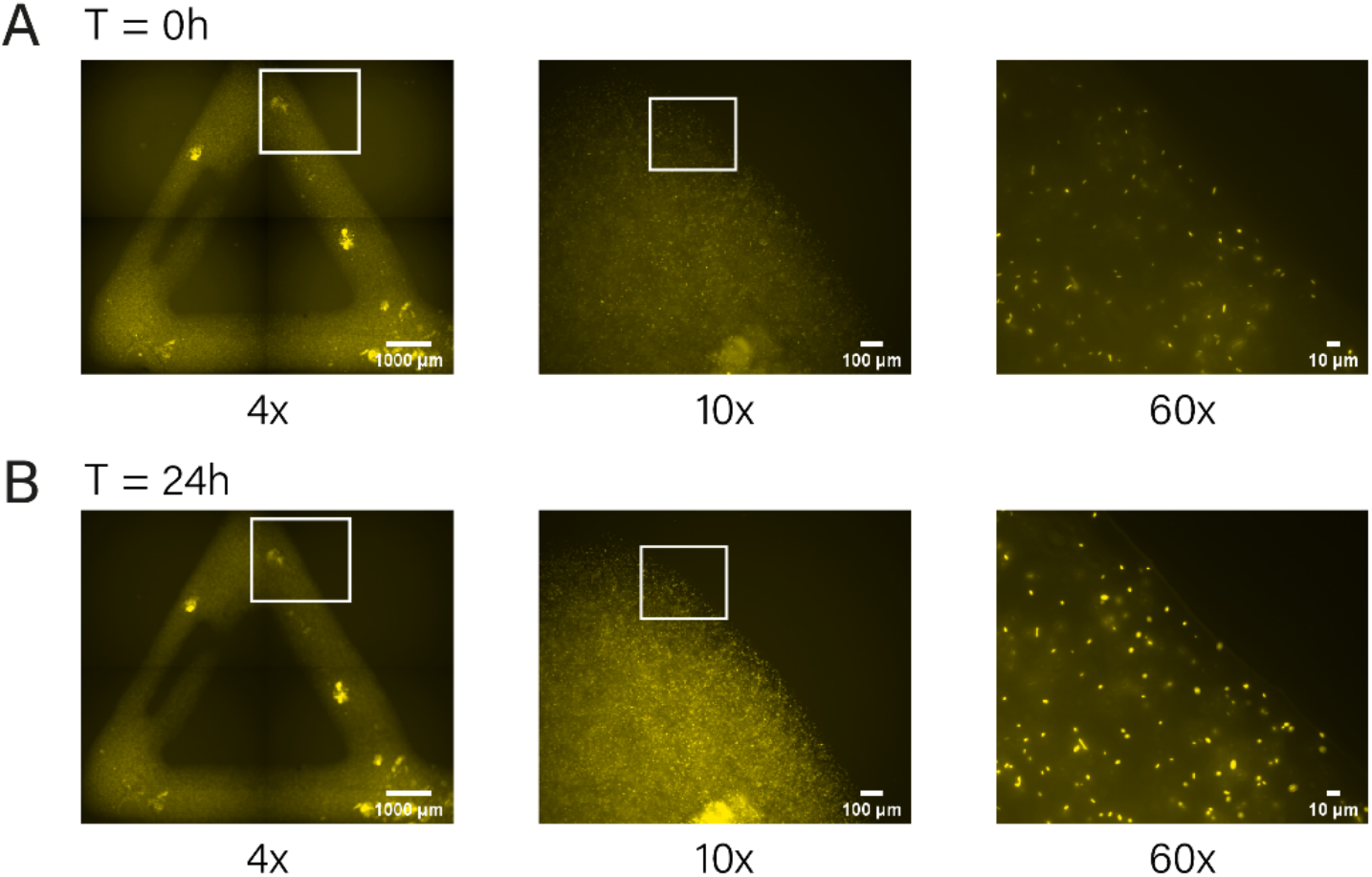
Printed triangular structure with immobilized pConst-mRFP *E. coli* bacteria. After 24h of incubation a clear increase in fluorescence can be observed. Direct comparison of the respective magnification shows a clear immobilization of the singular bacteria to their respective positions.

### 3.3 Colony Growth Data

Colony growth was observed by analysis of the colony size of four colonies per 1h time step. The colony sizes were averaged and fitted with an exponential function to determine the generation time. The colony shapes are assumed to be ellipsoids, the axes x, y, and z were determined and the number of cells was calculated from the colony volume and an assumed average *E. coli* cell size of 1.5 µm^3^. All triplicates showed generation times ranging from 50 min – 60 min.

**Figure S4:**
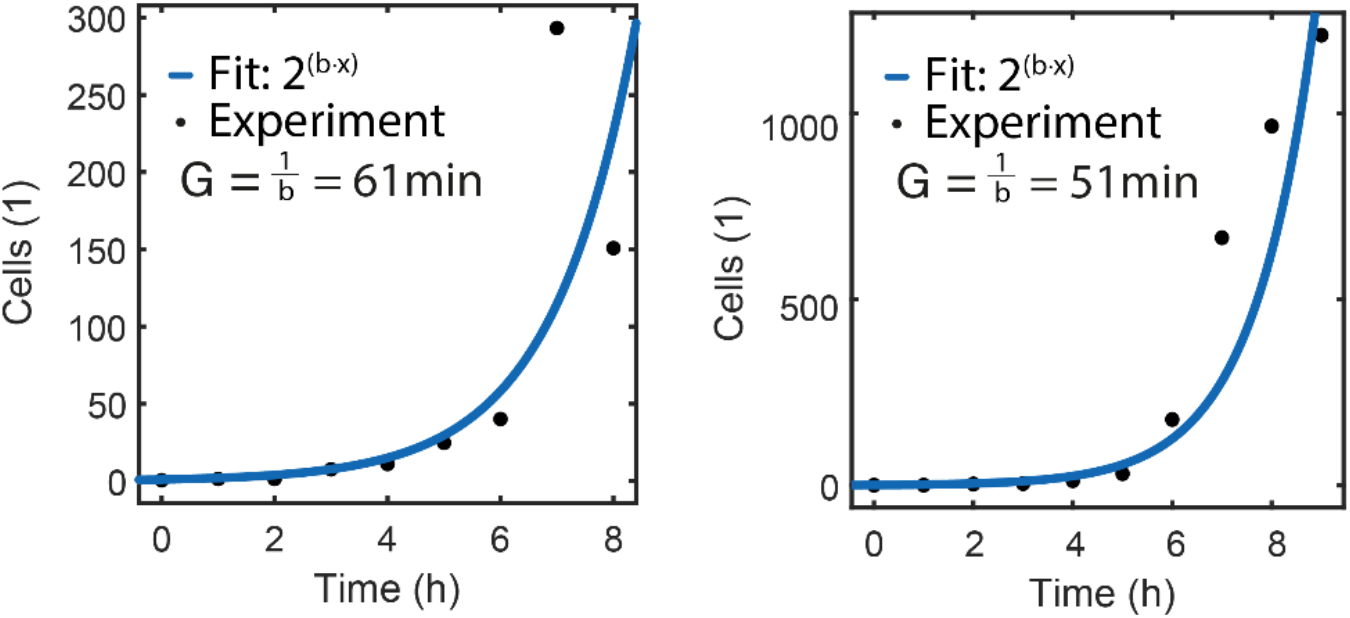
Data of two additional biological replicates of colony growth measurements over 0 h – 10 h in printed bac-ink. Together, the experiments resulted in a mean generation time of 55 ±6 min.

### 3.4 Gene Induction with aTc

For the experiments on spatiotemporal gene induction by diffusing aTc inducers, *E*.*coli* DH5αZ1 bacteria containing the plasmid pSB1A3-I13521 were used. For bulk characterization of the bacterial gene expression response (without bacteria-ink), measurements of 300 µL sample volumes were performed in a 96-well plate using a Fluostar Plate Reader. In the diffusion experiment shown in Fig. 3, a 2µl volume of bac-ink-aTc (500x) placed in the center of the print corresponds to a final concentration of 2.0 × aTc in the closed gel chamber, which expected to results in full induction of the bacteria.

**Figure S5:**
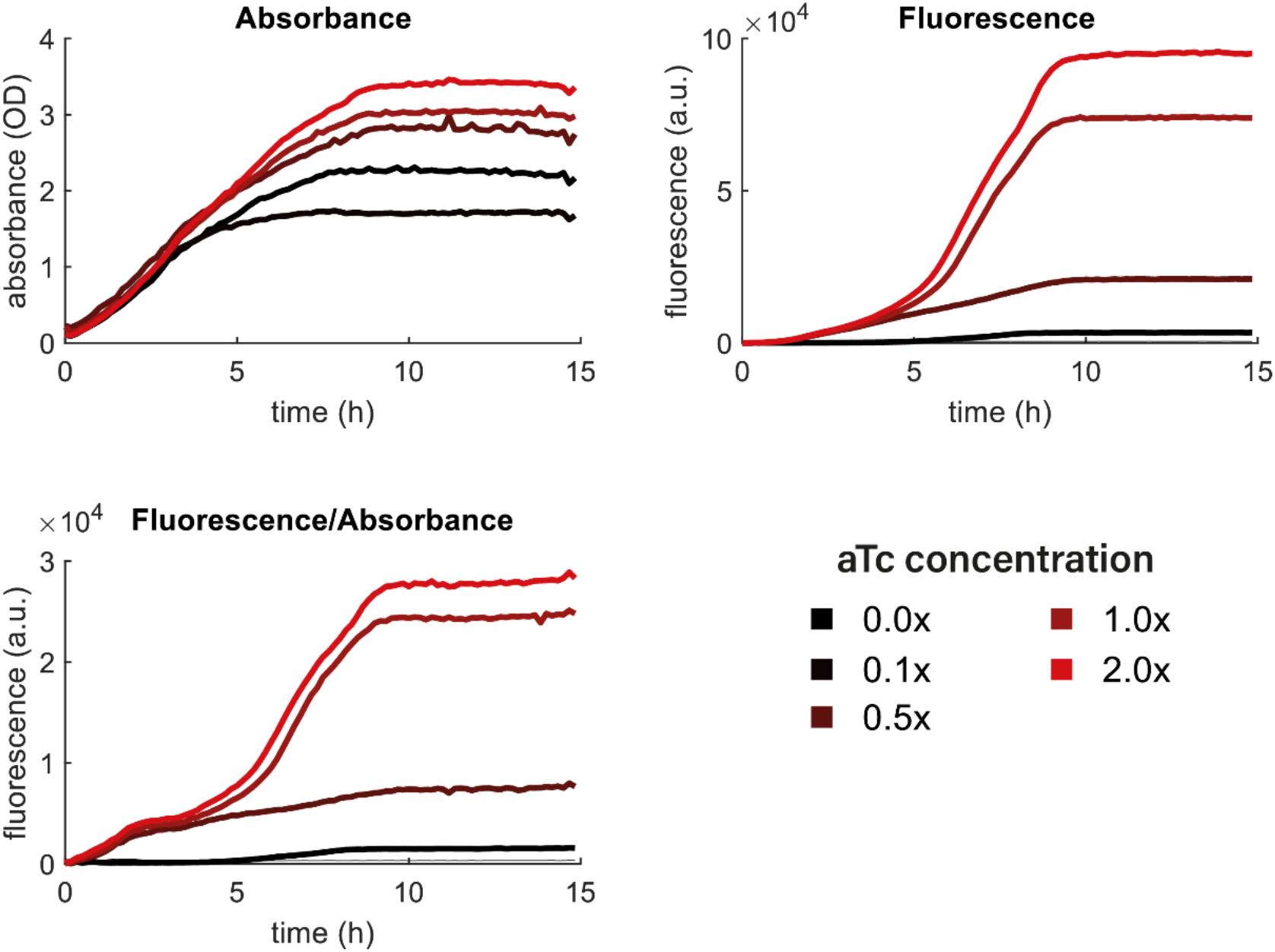
Absorbance, mRFP fluorescence and fluorescence/absorbance data for different aTc concentrations in bulk were determined in a plate reader experiment. A concentration of 1 × aTc corresponds to 0.1 µg/mL. In the diffusion experiment, a 2x (final) concentration is used, which is expected to result in full induction after equilibration of the inducer.

### 3.5 Gene Expression in Response to Diffusing aTc Inducers

**Figure S6:**
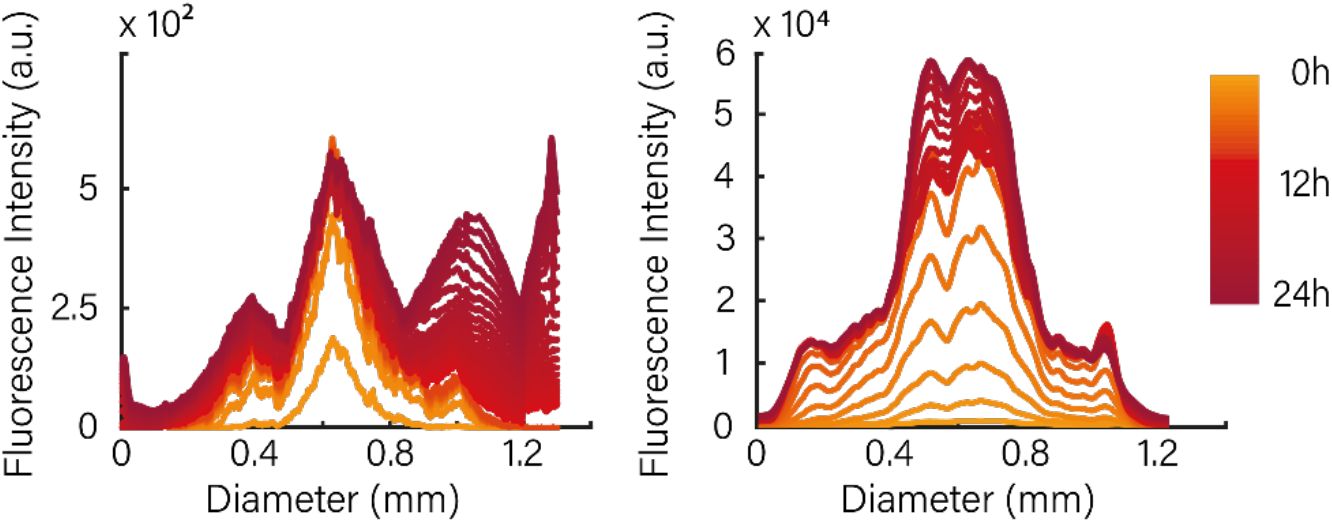
Spatiotemporal gene expression in response to aTc inducers diffusing from a spot in the center of a bioprint - data for additional biological replicates (cf. Figure 3B of the main paper). The substructure in the curves results from irregularities in the print and from stitching of the fluorescence micrographs.

### 3.6 Sender-Receiver Characterization

The sender-receiver system was tested in a bulk experiment and measured with a 96-well plate in a Fluostar Plate Reader with a total sample volume of 300 µl. In the absence of IPTG, receivers show a low fluorescence signal, which results from leaky AHL expression by the sendors. A correspondingly higher signal is obtained, when senders are induced with IPTG (1mM). Highest receiver levels are achieved by complete induction of the receiver cells with externally added AHL (200 nM). As negative controls, receiver cells without the addition of AHL or only sender cells were studied, which both showed no fluorescence.

**Figure S7:**
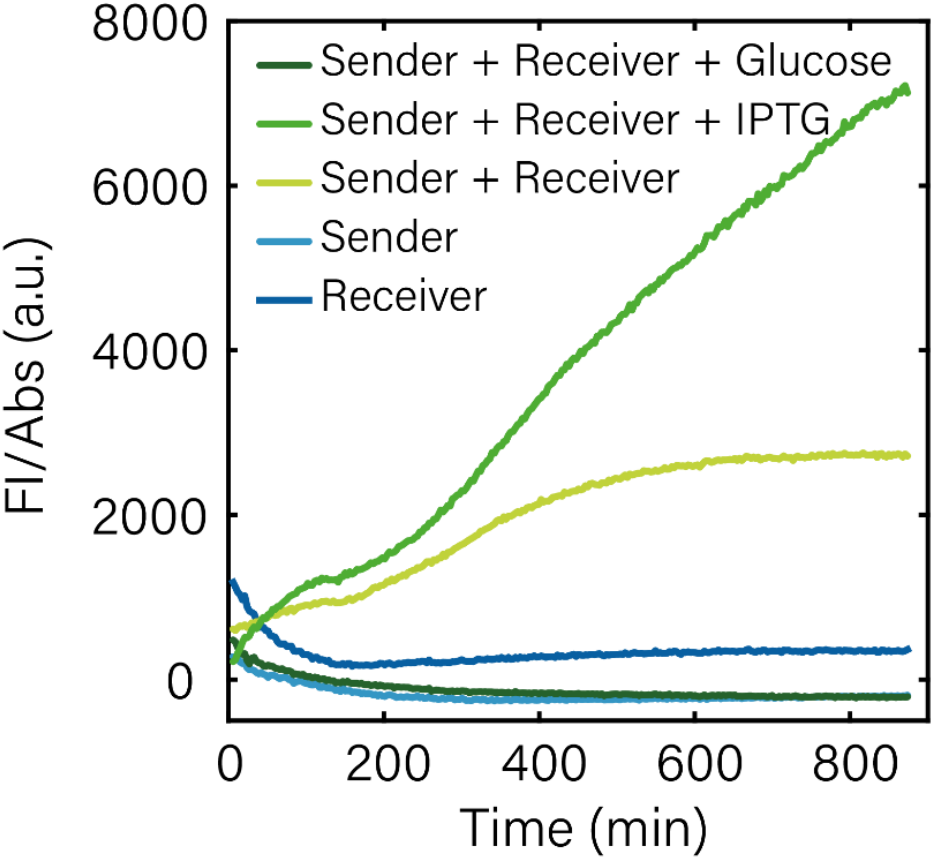
Sender-Receiver performance in plate reader measurements (Fluorescence/OD). GFP is generated only in the presence of senders and receivers. In the absence of IPTG, leaky expression of LuxI from the pLac promoter generates enough AHL to result in an appreciable GFP signal. Induction with 1 mM IPTG increases the signal. Leaky expression can be suppressed by the addition of glucose. Senders and receivers alone do not show any fluorescence in the GFP channel.

### 3.7 Fluorescence Data for the Bioprinted Sender-Receiver System

**Figure S8:**
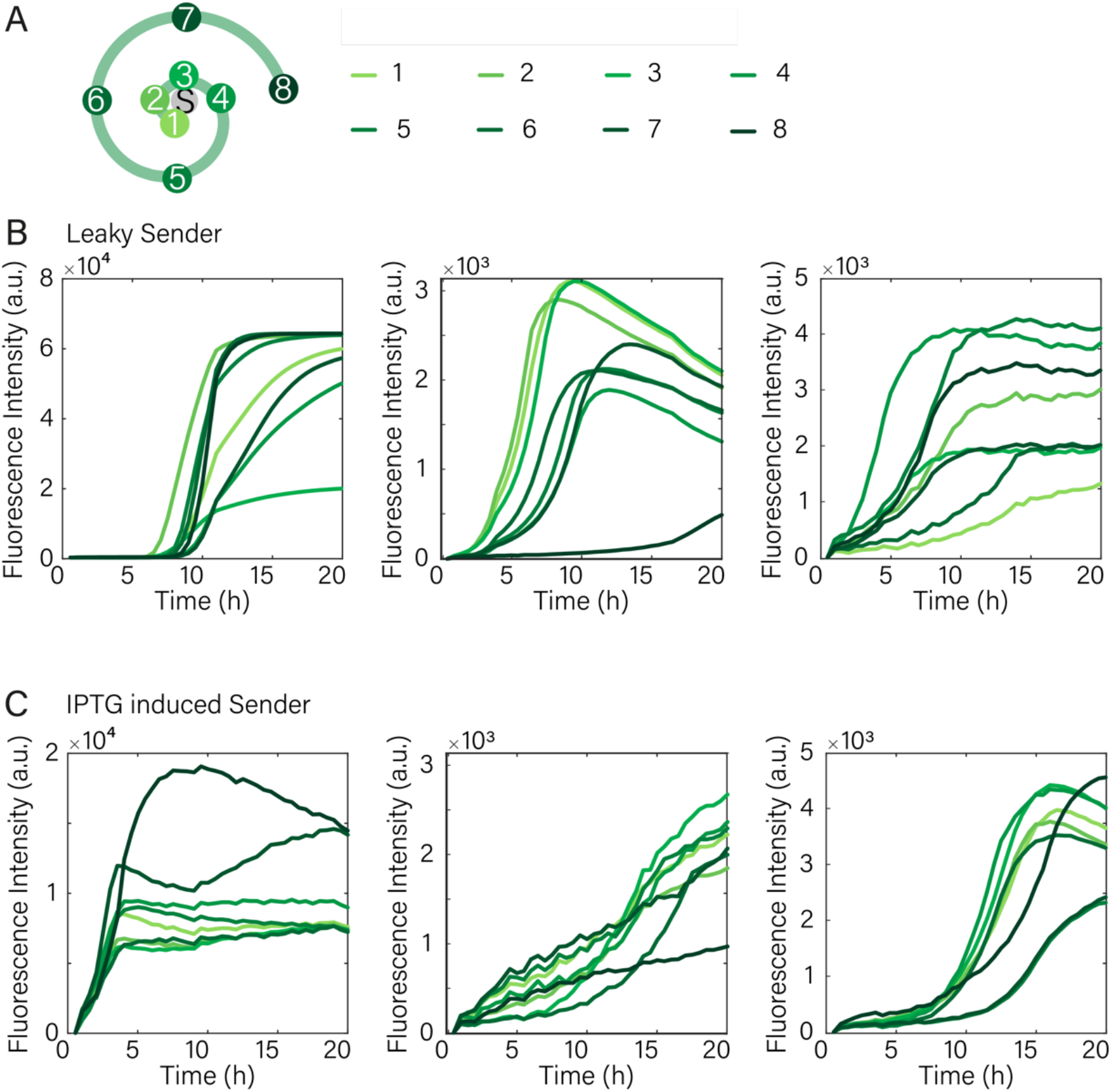
Fluorescence Data of triplicates with either leaky (low sender strength) and IPTG induced (full sender strength) sender cells (cf. Figure 3D).

### 3.7 Guiding Chemotaxis with Bacterial Boundaries

Experiments with printed *E. coli* boundaries were performed with pConst-mRFP bacteria, while printing with plain 1% alginate was used as a negative control. In all experiments the chemotactic cells were able to cover the complete agar surface over the course of 20h except for the majority of the enclosed area of the printed logo, when the logo was printed with non-motile bacteria. In the negative controls, the chemotactic bacteria were not affected by the printed structure and covered the complete plate (Figure S8).

**Figure S9:**
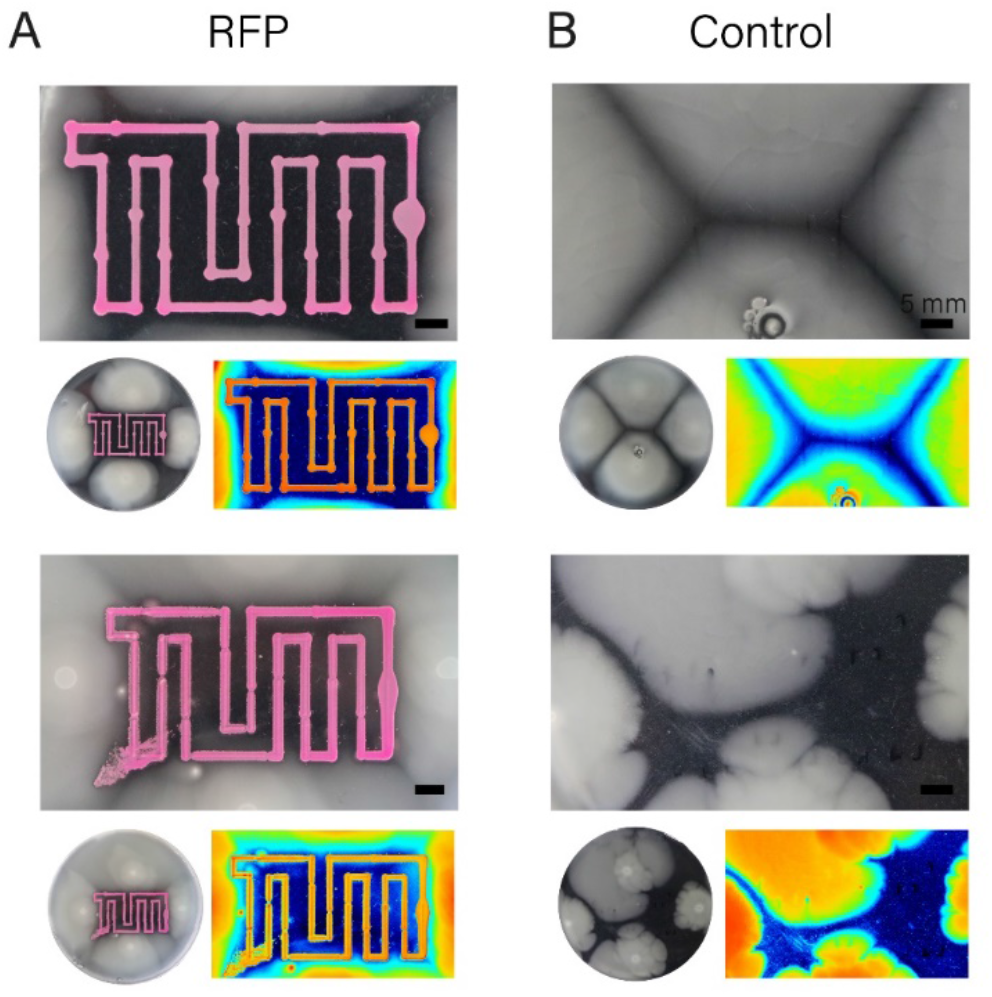
Experimental replicates for printed boundary experiments with (A) mRFP producing non-motile bacteria in the boundary and (B) with boundaries printed with alginate.

## 4. Computer Simulations

We modeled the spatiotemporal gene expression dynamics observed in the experiments discussed in Fig. 3 of the paper using a simple reaction-diffusion model. We considered the bioprint as a cylindrical gel slab with height *h* = 4.0 *mm* and radius *R* = 10 *mm*, where the diffusing inducer (or sender bacteria) is initially confined to the region *r* < *R*_0_. The inducer then diffuses isotropically within the gel with diffusion coefficient *D*. The bacteria are assumed grow exponentially with growth rate γ = *ln* 2 /*t*_*d*_, where *t*_*d*_ is the doubling time, consistent with the observations of Fig. 2A. Bacteria produce fluorescent proteins depending on the local inducer concentration, which is modeled with a Hill function with a threshold concentration of *K*, Hill coefficient *n*, and maximum production rate α. This leads to the following differential equations for inducer concentration *c*(*r, t*), bacteria number *N*(*r, t*) and total amount of protein *P*(*r, t*) (as the experimental observable is total fluorescence, we model the total protein number):

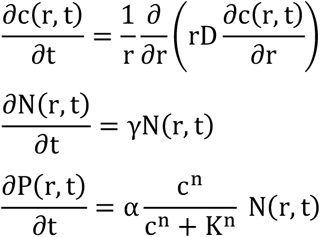

The equations were solved numerically using the Matlab pde solver *pdepe* with no-flux boundary conditions for *c*.

### 4.1 Diffusing Inducer

For the experiments with diffusing aTc, as an initial condition we used a 0.39 mM concentration of aTc confined to a cylinder with radius *R*_0_ = 0.4 *mm*. The other parameters were set to α =, *n* = 2, *K* = 40 *nM*. A screen of the diffusion coefficient showed the best match with our experimental data for a value of *D* = 200 μ*m*^2^/*s* (Fig. S10).

**Figure S10:**
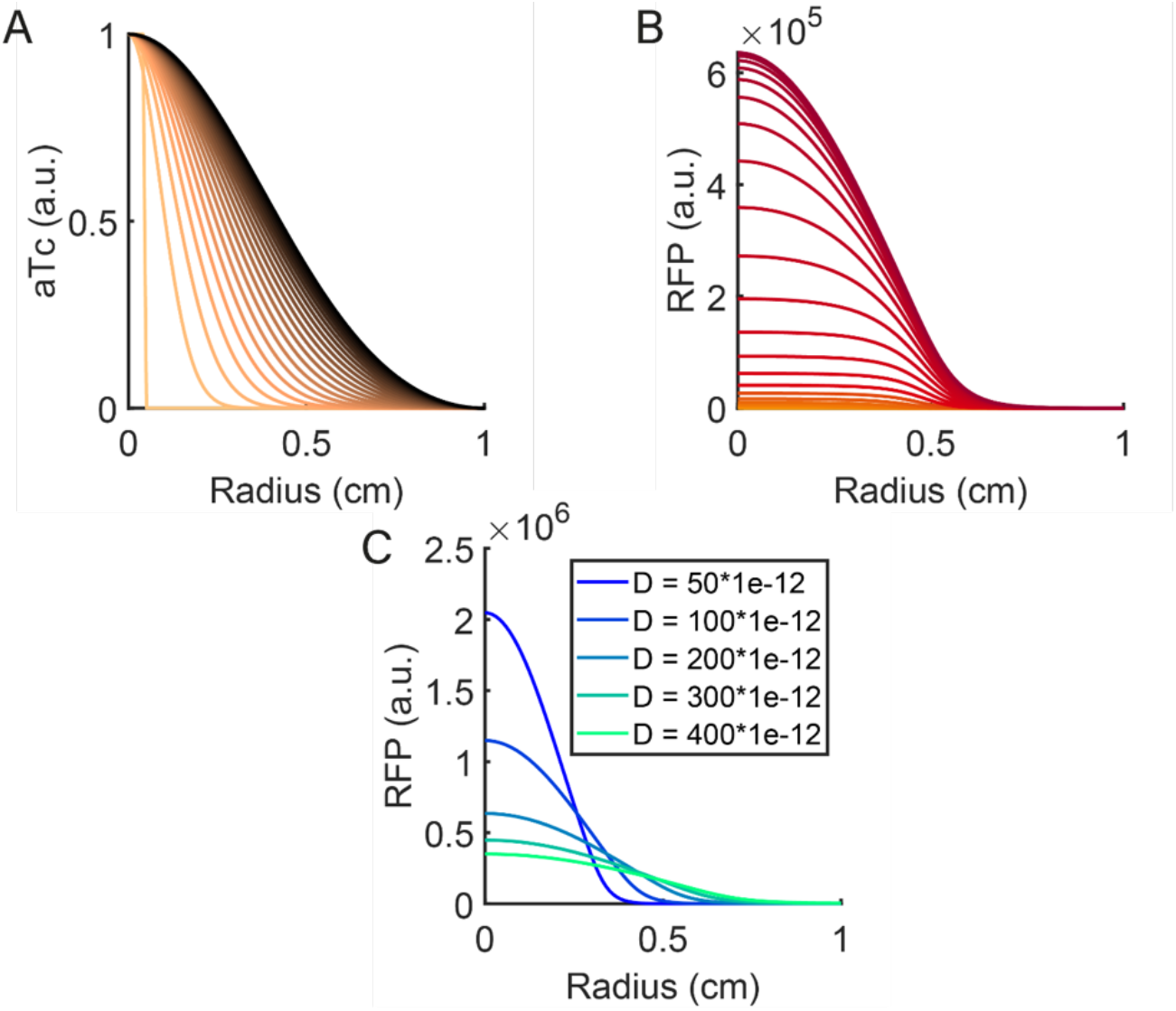
A) shows diffusion of aTc away from the center (r=0), while B) shows the expected RFP signal at different positions. The color gradients indicate the progress over time. C) shows how the aTc diffusion coefficient influences the RFP profile. The expected RFP intensity is depicted after 10h for diffusion coefficients ranging from 50 μm^2^/s to 400 μm^2^/s. Further model parameter values are *n* = 2 and *K* = 0.01495 µ*g*/*L*.

### 4.2 Sender-Receiver Simulations

The sender-receiver system requires a more complex model that incorporates growth of the senders and receivers, production of the signal AHL by the senders, and GFP expression by the receivers in response to AHL. In order to qualitatively capture the experimental observations made in Figs. 3D and S8, we also need to consider the degradation of AHL and GFP. Furthermore, the experiments indicate a time lag between growth/expression of the senders and growth/expression of the receivers. This parameter appears to vary between experiments and is expected to have a strong impact on the spatiotemporal dynamics of GFP expression. It is actually reasonable to assume such a lag as the receivers are printed in the experiments (and thus are exposed to a brief high temperature “shock”), while the senders are simply applied to the center of the printed to the structure.

- *Sender cells*: We assume that the senders are confined to the central region (r < *R*_0_ = 0.4 *mm*) of a cylindrical gel with *R* = 10 *mm*. Initially, a 2 μL volume of bacteria ink with cells at a concentration of ≈ 2 × 10^8^ cells/mL is applied to the center, i.e., the initial number of senders is 4 × 10^5^ sender cells. Based on bulk growth experiments, we expect the cell number to saturate at a ≈ 20 × higher value, i.e., *N*_*max*_ = 8 × 10^6^ cells.

For simplicity, we assume logistic growth of the senders, i.e.,

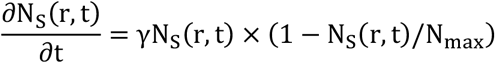

with Y as above.

#### Receiver cells

Receiver cells, which are only present for *r* > *R*_0_, are modeled just as the sender cells, but we now explicitly model a time lag via the sigmoidal function *s*(*t, t*_*lag*_): = 1/(1 + exp(™κ (t ™ t_*lag*_)), where κ is controls the smoothness of the transition to growth and is set to the (arbitrary) value 10^−3^. Hence,

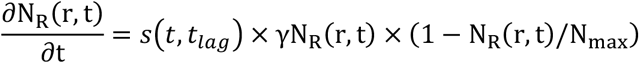

#### AHL turnover

We further assume steady state production of LuxI by the senders at a concentration of 1 μM (or a copy number of 1000). Each LuxI enzyme will generate AHL at a rate of ≈ 1 molecule/second. The contribution to the AHL production of each sender bacterium within the sender volume 2 μL is thus

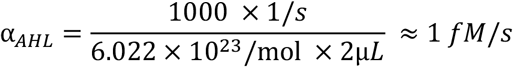

The overall reaction-diffusion dynamics of AHL is then given by the equation:

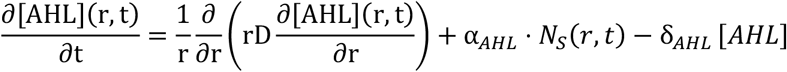

The degradation rate of AHL is set to δ_*AHL*_ = 0.5. 10^−5^*s*^−1^, while its diffusion coefficient is set to *D* = 1000 µ*m*^2^/s (for simulations with different diffusion coefficients see Fig. S12).

#### GFP production

Just as in the simple induction experiment (4.1), we assume the production of the total number of fluorescent proteins *N*_*GFP*_ to be proportional to the number of bacteria. Due to the longer run-time of the experiments, we explicitly account for a reduction of gene expression in the stationary phase (phenomenologically modeled via logistic growth as for the bacteria themselves). Further, we cannot neglect degradation of GFP, which we assume to occur roughly within 1 day, i.e., δ_*GFP*_ = 10^−5^/*s*:

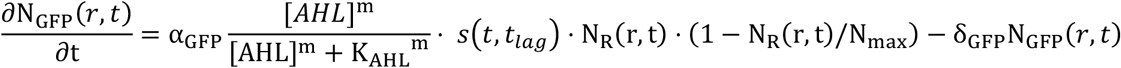

The Hill coefficient for AHL induction is set to *m* = 1, *K*_A*HL*_ = 10*nM*. For our purpose, the expression rate *α*_*GFP*_ is an arbitrary scaling parameter and is set to 1/s.

These equations can reproduce the various fluorescence time courses observed in the sender-receiver experiments quite well (cf. Fig. S11). Specifically, they capture the observation that longer lag times tend to sharpen the transition – at later times AHL has already distributed in the gel, and all bacteria are induced above threshold without any spatial dependence. The model further explains that at low sender strength and without time lag there is a stronger differentiation than at high sender strength without lag, or, alternatively, at low sender strength, but with time lag. The simulations also suggest that within one print bacteria at different positions might show slightly different growth behavior, leading to the variability observed in the curves displayed in Fig. S8.

**Figure S11.**
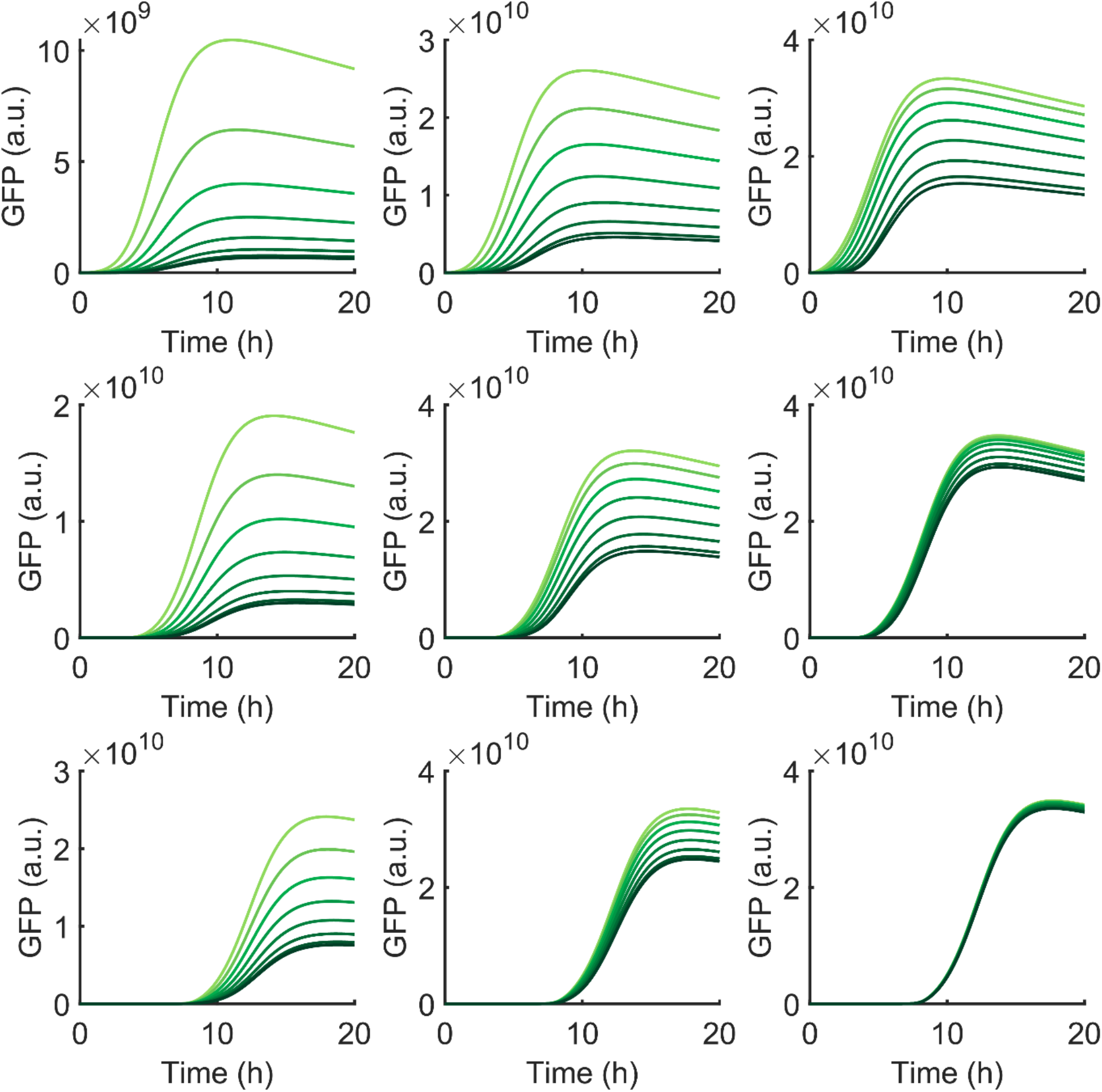
Simulated GFP intensities for the sender-receiver experiments. The response by the receiver bacteria is strongly dependent on sender strength and the time lag between sender and receiver growth. From left to right, the sender expression rate is varied uninduced to induced, i.e., *α*_A*HL*_ = 0.01,0.1, And 1 *fM*/*s*/*cell*. From top to bottom the lag time is varied from 0 over 4 h to 8 h. Both higher expression and larger lag time sharpen the response.

**Figure S12.**
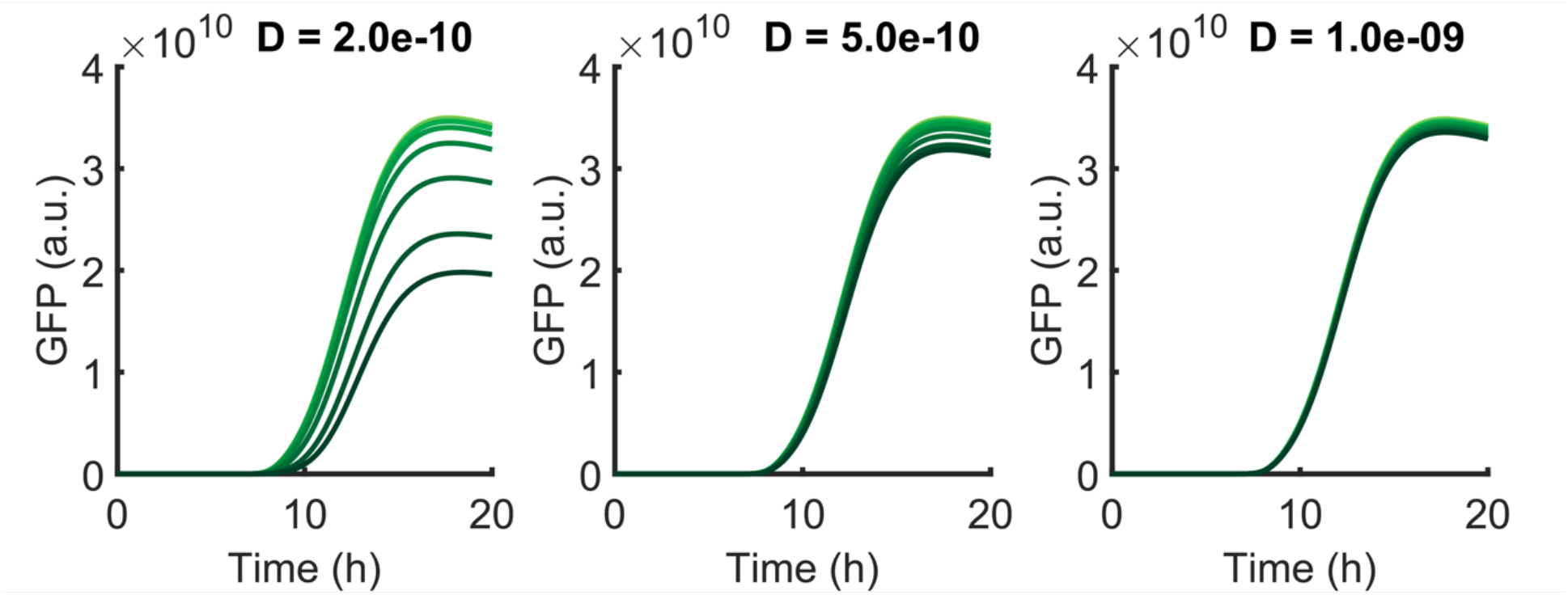
Influence of diffusion coefficient on sender-receiver dynamics for a lag time of 8 hours. Faster diffusion has a sharpening effect (D is given in units of *m*^2^/*s*) similar to lag time and expression strength (see Fig. S11).

