## Supplementary figures and images for "Bacterial growth, communication and guided chemotaxis in 3D bioprinted hydrogel environments"

### Supporting Video

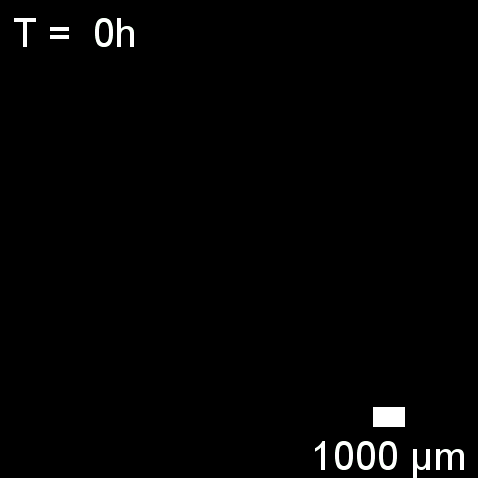
